# Neural Gain Modulation Propagates from Posterior to Anterior Brain Regions to Optimize Orientation Perception in Chronic Astigmatism

**DOI:** 10.1101/2025.04.13.648557

**Authors:** Sangkyu Son, Hyungoo Kang, HyungGoo R. Kim, Won Mok Shim, Joonyeol Lee

## Abstract

While visual impairments commonly occur daily, many individuals fail to recognize these distortions. Yet, the brain’s role in adapting to distorted sensory inputs remains largely unknown. In this study, we focused on how the brain recalibrates physical orientation-specific blur after chronic exposure to astigmatism. By reconstructing the population orientation tuning response from electroencephalogram activity patterns and estimating neural gain modulation using an optics-based computational model (*data from 42 participants, including 15 females*), we found enhanced neural gain for underrepresented orientations and reduced gain for overrepresented ones, especially in individuals with long-term astigmatism. The strength of the gain modulation correlated with the optimization of orientation perception in these participants. Furthermore, this *push-pull* neural gain modulation dynamically propagated from the posterior brain regions to others, and the strength of the propagation correlated with the degree of perceptual optimization. In contrast, short-term exposure resulted in transient and short-lived neural optimization, characterized by a relatively stronger anterior-to-posterior transference pattern. These results show how feature-specific information is modified across the entire brain in response to systematic visual distortion, revealing duration-dependent strategies the brain employs to handle sensory impairments.

## INTRODUCTION

Optical lenses, susceptible to imperfections from external forces and scratches, have counterparts in modern digital cameras that use sophisticated algorithms for image optimization, including selective pixel masking and contrast enhancement. Similarly, the human visual system—comprising the eye and brain—exhibits a remarkable capacity for compensatory adjustment in response to visual distortions, much like photo editing software corrects image flaws. This adaptive process allows for improved visual discrimination in the face of refractive errors such as myopia, hyperopia (Radhakrishnan et al., 2015; Webster et al., 2002), keratoconus (Barbot, Park, et al., 2021), and astigmatism (Ohlendorf et al., 2011; Vinas et al., 2012).

Astigmatism, in particular, is a widespread condition, affecting a significant portion of the adult population (up to 40 – 60%) (Hashemi et al., 2018; Williams et al., 2015; Wong et al., 2000) and notably diminishing key visual functions (Atchison & Mathur, 2011; Kobashi et al., 2012; Wolffsohn et al., 2011), such as orientation perception and the corresponding neural representation in the visual cortex (Son et al., 2021). This impairment stems from the deviation of the cornea and lens from their ideal symmetrical shape, resulting in a distortion akin to viewing through a rugby-ball-shaped lens. Notably, perceptual enhancements in astigmatism are tilt-dependent (Georgeson & Sullivan, 1975; Sawides et al., 2010; Son et al., 2022), indicating a need for orientation-specific interventions. Therefore, the improvement in vision among astigmatic individuals is attributed to cerebral rather than ocular adjustments, as the eye alone is incapable of multi-focal accommodation inherent to astigmatism (compare red and gray solid lines in Figure 1A). The cerebral contribution to restoring orientation perception in astigmatic vision is also exemplified by often reported side effects when the full optical correction was done purely based on optical aberration without considering extra-retinal factors (Mitchell et al., 1973; Son et al., 2022).

**Figure 1.**
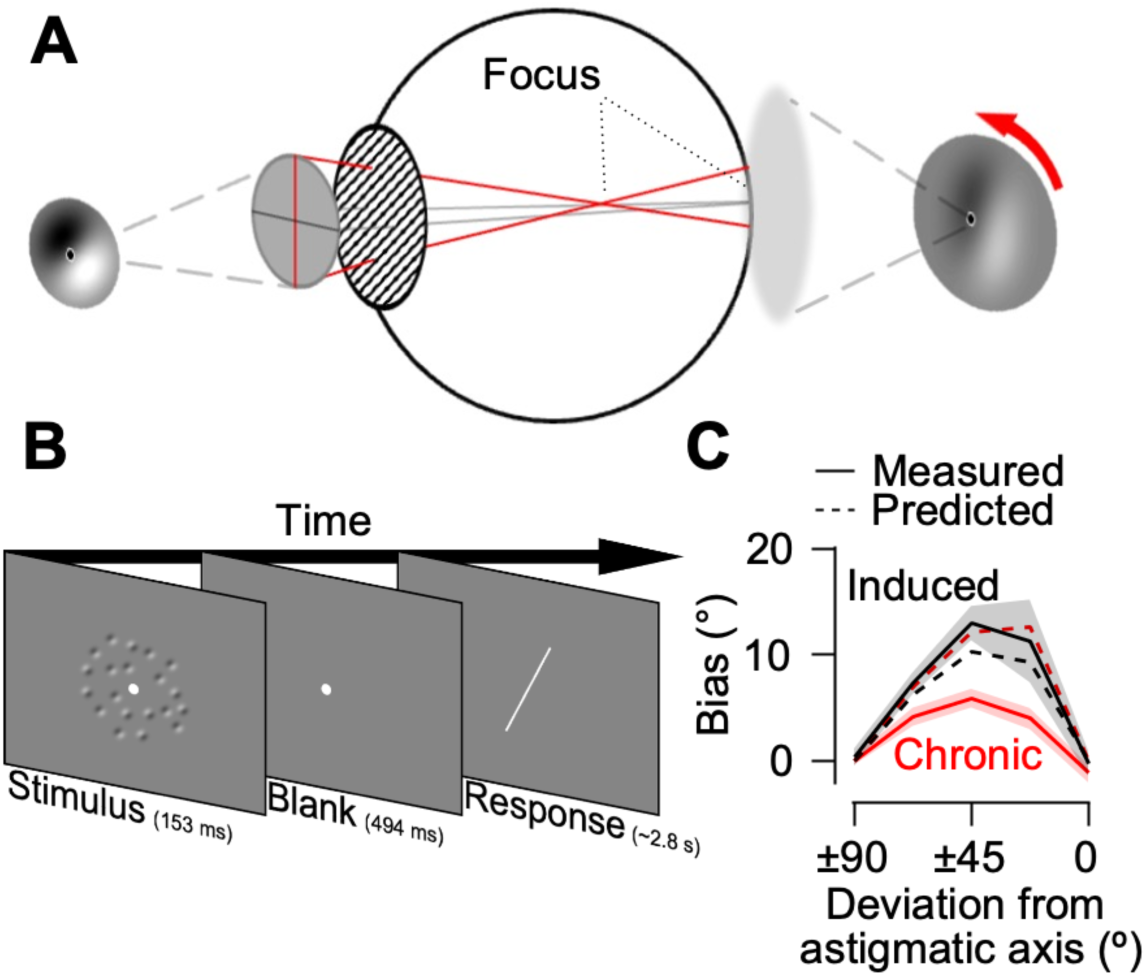
Experimental design and effect of astigmatism on orientation perception. **(A)** Impact of astigmatic optical deformation on orientation information. Light rays passing through an angle perpendicular to the astigmatic axis undergo greater refraction and focus before reaching the retina (red lines). This results in an elliptical optical blur, causing the orientation of the Gabor stimulus on the retina to appear tilted away from the astigmatic axis (red arrow). Astigmatic vision was simulated either through the participant’s own chronic astigmatism (referred to as *chronic*) or by experimentally inducing an astigmatic refractive error in individuals with normal vision (referred to as *induced*). **(B)** Participants were presented with randomly tilted Gabor stimuli at the fovea. After a brief presentation, they reported the perceived mean orientation by adjusting an orientation bar. **(C)** Perceptual biases of the chronic group (solid red lines), the induced group (solid black lines), and the optics-based model prediction (dashed lines) as a function of deviation from the astigmatic axis. The shaded regions represent ±1 SEM.

The role of neural modulation at the early stages of visual processing is crucial in this context, as the rectification of astigmatic distortion necessitates extracting and processing orientation information within the visual pathway before any higher-level visual processing can occur. Despite this understanding, direct evidence elucidating the precise mechanisms by which the brain alters sensory input to manage astigmatism effectively remains sparse. This paper investigates the neural strategies that enable individuals with astigmatism to perceive their environment with enhanced clarity.

To understand the temporal evolution of neural and perceptual compensation in astigmatic vision as a function of exposure duration, we analyzed multivariate electroencephalogram (EEG) responses to orientation stimuli in participants with chronic astigmatism and compared them to responses from participants with normal vision who were subject to acutely induced astigmatism. Through a model-based EEG analysis, we found that chronic exposure to astigmatic vision caused astigmatic axis-specific push-pull gain modulation of EEG responses in the visual cortical regions, which propagated to other brain regions and contributed to the restoration of orientation perception. Short-term exposure to astigmatism resulted in only transient and weak gain modulation with an anterior-to-posterior propagation pattern, which did not correlate with orientation perception.

## RESULTS

Forty-two participants reported the perceived orientation of Gabor stimuli under the influence of astigmatic vision (Figure 1B). We recruited two groups of participants based on the innate optical properties of each eye, which were pre-identified by an auto-refractometer (Huvitz, HRK-7000). One group of participants had chronic astigmatism and the other had near-perfect vision. The group with chronic astigmatism (chronic astigmatism group) performed the task mainly based on their own eye’s cylindrical refractive error (−2.36±0.23 diopter; see Table 1 for details), while the group with the near-perfect vision performed the task with +2.00 diopter cylindrical lens placed in front of their healthy eye (induced astigmatism group). The spherical refractive error, accommodation, and influence of the other eye were controlled in both groups of participants. We compared the neural orientation representations of the two groups using 64-channel EEG recordings.

**Table 1.** Measured refractive errors of the chronic astigmatism group.

| # | Sex | Age | Which eye | Spherical (D) | Cylindrical (D) | Axis (°) |
| --- | --- | --- | --- | --- | --- | --- |
| 1 | Male | 19 | Left | -6.50 | -1.25 | 160 |
| 2 | Male | 23 | Left | -5.00 | -3.00 | 175 |
| 3 | Male | 19 | Right | -17.00 | -1.00 | 175 |
| 4 | Female | 19 | Left | -5.25 | -2.25 | 170 |
| 5 | Female | 19 | Right | -7.25 | -2.25 | 5 |
| 6 | Male | 25 | Left | -2.00 | -1.50 | 170 |
| 7 | Male | 32 | Right | -9.75 | -1.25 | 5 |
| 8 | Male | 19 | Right | -4.50 | -2.75 | 170 |
| 9 | Female | 20 | Right | -5.25 | -2.00 | 5 |
| 10 | Male | 23 | Left | -6.00 | -3.25 | 170 |
| 11 | Female | 19 | Right | -6.75 | -3.25 | 175 |
| 12 | Female | 18 | Left | -6.50 | -3.75 | 0 |
| 13 | Male | 22 | Right | -3.75 | -1.00 | 0 |
| 14 | Male | 22 | Right | -6.00 | -2.50 | 170 |
| 15 | Female | 20 | Left | 0.00 | -2.50 | 175 |
| 16 | Female | 20 | Right | -3.75 | -1.25 | 5 |
| 17 | Male | 28 | Left | -3.75 | -2.25 | 163 |
| 18 | Male | 19 | Left | -8.50 | -1.00 | 170 |
| 19 | Male | 26 | Left | -3.00 | -4.00 | 0 |
| 20 | Female | 19 | Left | -5.25 | -1.00 | 170 |
| 21 | Male | 24 | Left | -5.75 | -3.25 | 170 |
| 22 | Male | 24 | Right | -3.00 | -5.00 | 0 |
| 23 | Female | 19 | Right | -5.25 | -3.00 | 10 |

Note, D and ° denote diopter and degree, respectively.

### Chronic exposure to astigmatic vision alters neural orientation tuning responses

Optical deformation caused by astigmatism induces biases in the physical orientation of the retinal image of the Gabor stimulus when its orientation is oblique to the astigmatic axis (Figure 1A; see Supplementary Figure 1A for a simulated illustration of retinal images in astigmatism). If perception mirrors the deformed retinal image, the perceived orientation should match the predicted orientation calculated from the optics-based model that reflects the optical property (cylindrical refractive error) of astigmatic vision. In the induced astigmatism group, perceived orientation indeed matched the prediction of the optics-based model (Figure 1C; compare black solid and dashed line; t_(18)_ = 0.896, p > 0.05). However, in the chronic astigmatism group, the perceptual biases were much smaller than those predicted from the optics-based model (comparison between red solid and dashed lines; t_(22)_ = −3.244, p = 0.004), and smaller than those observed in the induced astigmatism group (comparison between the black and red solid lines; t_(40)_ = −2.695, p = 0.010), as reported previously (Son et al., 2022). This contrast was also observed when we predicted perceptual biases caused by optical deformation using ISETBIO (Cottaris et al., 2019), an open-source tool for advanced optics modeling. This suggests that the observed differences were unlikely to result from other factors, such as image magnification caused by the lens. Moreover, the perceptual bias predicted by our simpler optics-based model closely matched that of the ISETBIO simulation (Supplementary Figure 1B).

Simultaneously recorded EEG activity showed differences in the amplitude of the event-related potentials (ERP). The P1 component of the ERP was smaller in the astigmatic vision condition than in the emmetropic vision condition (where retinal distortion was corrected, Supplementary Figure 3A), consistent with previous reports (Regan, 1973; Sokol, 1983; Son et al., 2021). This is mainly due to the immediate contrast changes in retinal images caused by the axis-dependent blur of visual input in astigmatic vision. However, further examination of ERP amplitude variations as a function of stimulus orientation did not reveal strong systematic changes with respect to the astigmatic axis. Although reduced ERP responses were observed for orientations near the astigmatic axis in some channels, none of these effects reached statistical significance. Additionally, there was no clear evidence of compensatory responses for optical blur, such as orientation-specific changes in ERP amplitude, particularly in the chronic astigmatism group (Supplementary Figure 3B).

Therefore, we adopted multivariate approaches to extract visual orientation information from the EEG activity patterns, enabling a more robust and sensitive measure of orientation representation. Specifically, we derived orientation tuning functions by computing Mahalanobis distances (see *Methods* for details), to assess whether the strength of orientation information in the retinal inputs was indeed reduced due to optical blur near the astigmatic axis (Figure 2A, left panels). If the retinal inputs are the only source of the orientation tuning function estimated from the EEG activity pattern, the neural discriminability of stimulus orientations near the astigmatic axis should decrease (probably due to selective response reduction of the neurons whose preferred orientations are aligned with the astigmatic axis), and the tuning response profile should be shifted away from the astigmatic axis (Figure 2A, second right panel). However, when we estimated the orientation tuning functions from the EEG activity measured from posterior electrodes in the chronic astigmatism group, the tuning was not skewed away from the astigmatic axis. Rather, the presented stimulus orientation was correctly encoded in the EEG activity for an extended period.

**Figure 2.**
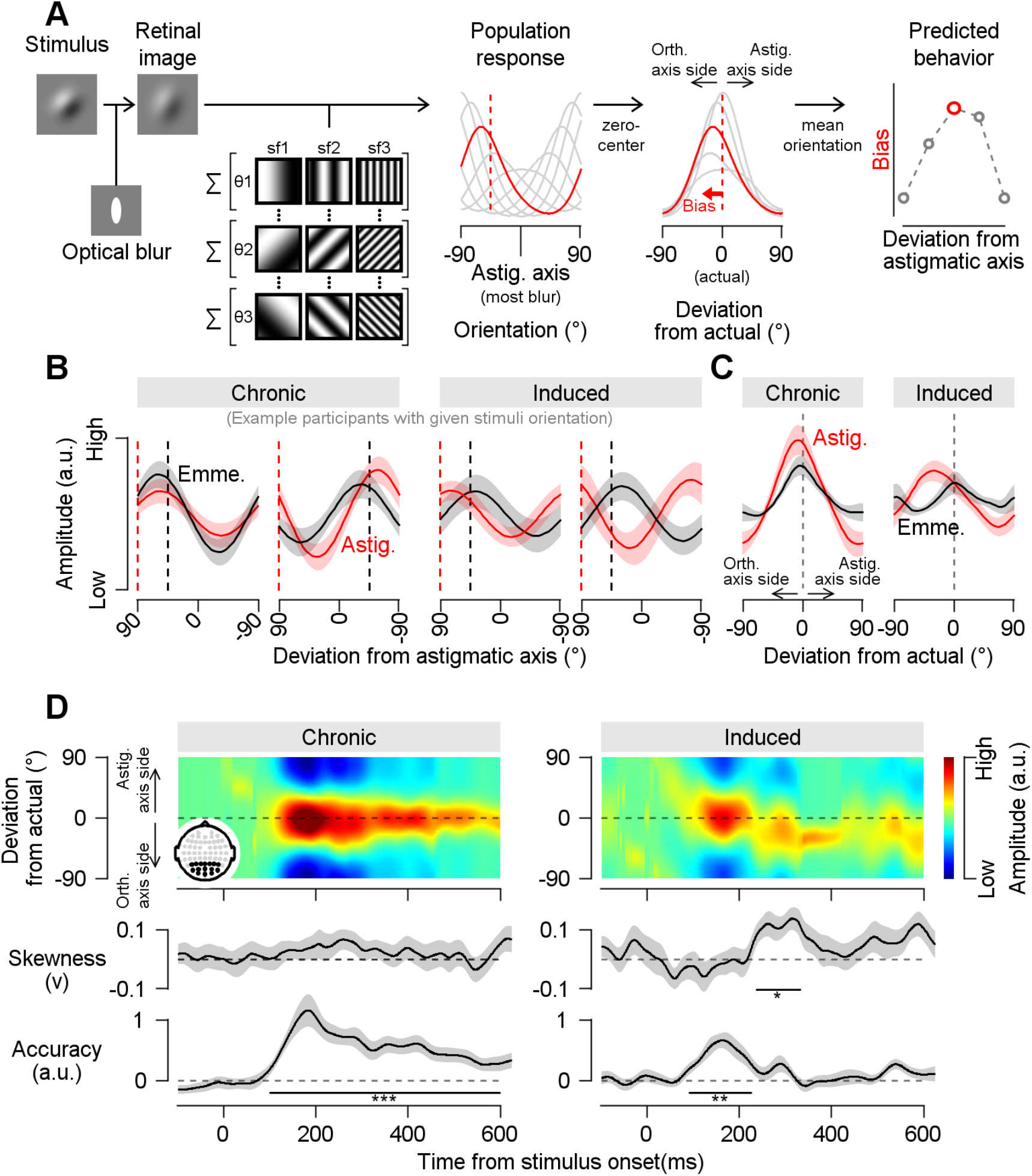
Restoration of the neural orientation tuning responses in chronic astigmatic vision. **(A)** Optics-based model of astigmatic vision. First column: The optically blurred retinal image in astigmatic vision was simulated by convolving visual stimulus with an elliptical-shaped kernel. The ellipticity of the kernel was estimated based on each participant’s cylindrical refractive errors. From the retinal image, population orientation tuning responses were estimated from a composite of hypothetical neurons with varying orientation and spatial frequency preferences. Second column: The population tuning responses to the presented stimulus orientation (−45°) are shown as a red solid line. The dashed red line represents the presented stimulus orientation, while tuning responses to other orientations are shown as gray solid lines. Third column: The predicted response to the presented orientation is centered at 0°, while responses to orientations near the orthogonal axis and the astigmatic axis are positioned to the left and right, respectively. Fourth column: The predicted behavioral responses were estimated from the mean orientation of the model’s population responses. Behavioral bias was calculated as the difference between the presented stimulus orientation and the model prediction. The red circle indicates the predicted bias for the example stimulus (−45°), while gray circles represent predicted biases for other stimulus orientations. **(B)** Orientation tuning responses measured between 250 and 350 ms after stimulus onset. Data from four example participants are shown, with the two participants on the left belonging to the chronic group and the two on the right belonging to the induced group. Orientation tuning responses to the same oblique orientation stimulus are shown under both astigmatic (red) and emmetropic (black) vision conditions. The red dashed vertical lines indicate the orientation orthogonal to the astigmatic axis, where optical blur is expected to introduce a bias, while the black dashed vertical lines represent the actual stimulus orientation. **(C)** Average orientation tuning responses across participants between 250 and 350 ms relative to stimulus onset. Tuning responses to all oblique orientations relative to the astigmatic axis were realigned such that the response to the actual stimulus is centered, with the orientation orthogonal to the astigmatic axis positioned on the left side. **(D)** Orientation tuning responses to oblique orientations (relative to the astigmatic axis) as a function of time in the chronic (left) and induced (right) groups. The inset topography denotes the posterior electrodes used in the analysis. The subfigures below show skewness and the strength of the orientation response in the tuning. Von Mises functions were fitted for visualization purposes only. All shaded regions represent ±1 SEM, and stars denote statistical significance (*, p < 0.05; **, p < 0.01; ***, p < 0.001).

For example, in Figure 2B, the orientation tuning functions of representative participants in the induced astigmatism group—estimated between 250–350 ms after stimulus onset—show a shift away from the astigmatic axis under the astigmatic condition, compared to the emmetropic condition (dotted black lines indicate the orientation of the presented visual stimuli). In contrast, the peak values of the orientation tuning functions in the chronic astigmatism group were accurately aligned with the presented stimulus orientation under both vision conditions. This pattern was consistent at the group level as well (Figure 2C), with the chronic astigmatism group displaying accurate and unskewed orientation tuning, unlike the induced group. We note that the orientation tuning functions shown in the figures were generated by fitting the raw tuning responses—calculated using Mahalanobis distance—with a Von Mises function for improved visual clarity (See *Methods* for detail).

Furthermore, this accurate encoding of the presented stimulus orientation under the astigmatic conditions in the chronic astigmatism group was maintained for extended periods (Figure 2D, left panel; cluster-based permutation test (Maris & Oostenveld, 2007); skewness, p > 0.05 for whole duration; accuracy, p < 0.001 from 99 to 600 ms). In contrast, the orientation tuning in the induced astigmatism group showed a skewed tuning profile, with a transient accurate orientation encoding matching the stimulus orientation (Figure 2D, right panel; cluster-based permutation test; skewness, p = 0.015 from 236 to 334 ms; accuracy, p = 0.002 from 91 to 228 ms).

It is particularly intriguing to observe that the neural responses in the chronic astigmatism group did not match with the prediction of the optic-based model, although the orientation information in the retinal inputs is relatively similar in the chronic and induced astigmatism groups; the measured refractive errors were similar in two groups (t_(40)_ = 1.409, p > 0.05). The ability to encode stimulus orientation was comparable between the groups when participants’ eyes were fully corrected (Supplementary Figure 4; cluster-based permutation test; estimated orientation accuracy of chronic astigmatism group, p = 0.035 from 262 to 455 ms; estimated orientation accuracy of induced astigmatism group, ps < 0.003 from 79 to 247 ms and from 363 to 523 ms; skewness for both groups, ps > 0.05 for the whole duration). Therefore, the deviation of the orientation tuning function from the optics-based model prediction under astigmatic vision in the chronic astigmatism group cannot be explained by the difference in retinal inputs or orientation encodability. Then, the remarkable restoration of orientation perception and neural orientation decodability should have originated from extra-retinal information processes in the brain.

### Restoration of orientation encoding via neural gain modulation

To quantify the extra-retinal information processes that compensate for the retinal distortion of orientation information in chronic astigmatism, we built a model by further developing the optics-based model. Under the assumption that extra-retinal information processes systematically counteracted the distorted retinal input (Figure 3A, left panel), we introduced a push-pull component to the optics-based model that enhances the gain of orientation tuning responses near the astigmatic axis while reducing the gain near the orthogonal axis (Figure 3A, right panel). Functionally, the astigmatism-induced optical blur causes the orientation tuning responses to be shifted away from the astigmatism axis, and the extra-retinal push-pull gain mechanisms restore the shifted tuning responses to represent the physical stimulus orientations correctly. To test the validity of this model, we first estimated the best optics-based model that explained the measured orientation tuning responses from multivariate EEG recordings using a generalized linear model (GLM). Then we estimated the push-pull gain modulation that explained the remaining residual tuning responses (see *Methods* for detail). We estimated the optics-based model first, instead of estimating a combined, full model (optics-based model + push-pull gain modulation), to fairly compare how much of the EEG responses were shaped by the distorted retinal inputs in both groups.

**Figure 3.**
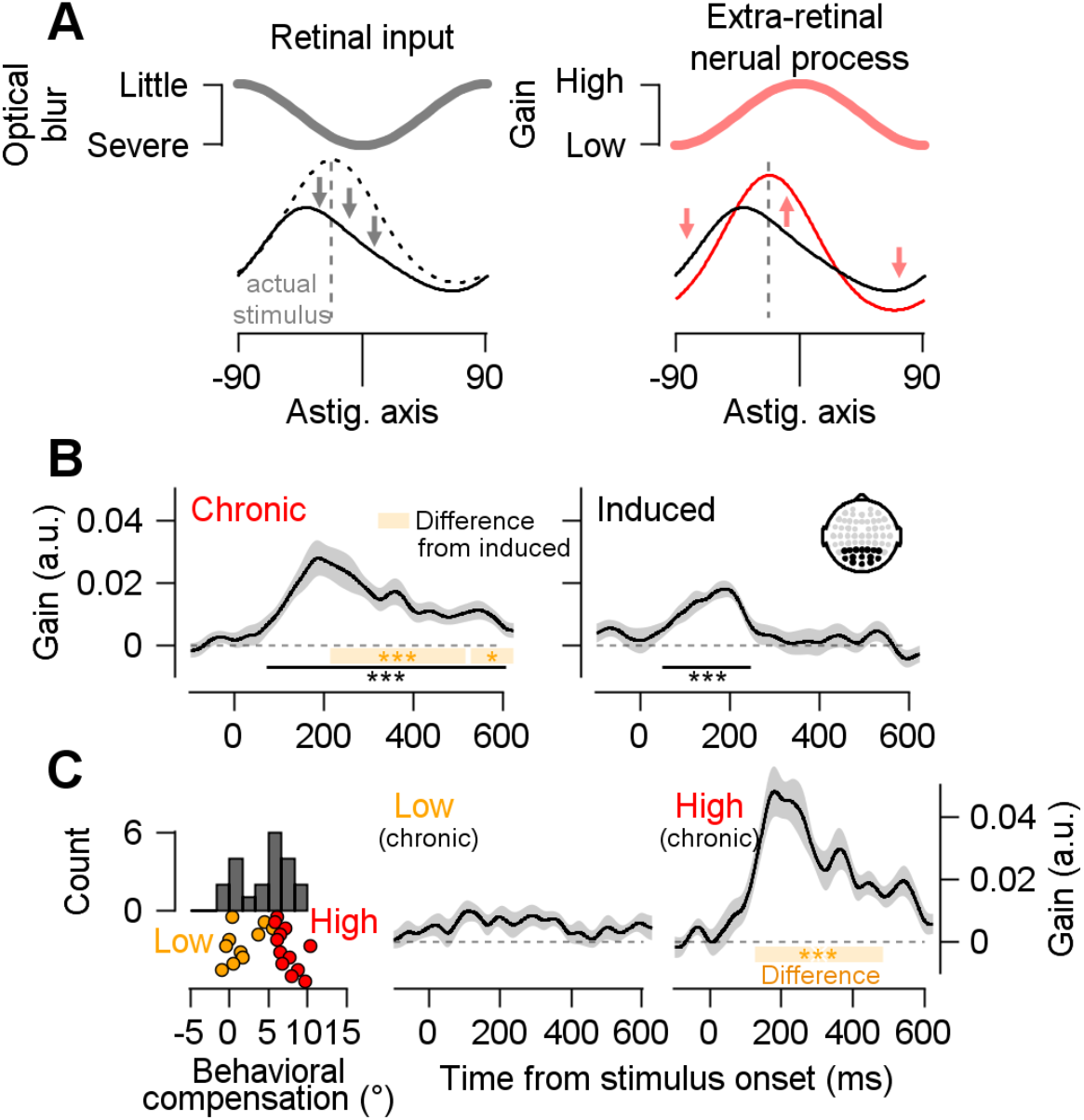
Neural gain modulation restores the orientation perception in astigmatic vision. **(A)** Left: The retinal input is degraded around the astigmatic axis, causing distortion in the orientation tuning response. As a result, the tuning function for the stimulus orientation (dashed line) becomes skewed (solid line). Right: If the extra-retinal neural process selectively increases the gain of neural responses to orientations near the astigmatic axis while decreasing the gain for orientations orthogonal to it, the skewed tuning response (black line; identical to the one on the left) can be restored to a more symmetrical shape (red line), effectively reversing the effects of optical blur. **(B)** Quantification of gain modulation estimated from the model shown in Figure 2. The orange shading represents the comparison between the chronic and induced groups. The inset topography indicates the posterior electrodes used in the analysis. **(C)** Temporal evolution of gain modulation in the chronic astigmatism group, comparing participants with high behavioral compensation to those low behavioral compensation. All shaded regions represent ±1 SEM. The number and color of stars indicate statistical significance (*, p < 0.05; **, p < 0.01; ***, p <0.001), with black denoting a one-sample t-test from zero and yellow denoting a paired t-test across groups.

The contribution of push-pull gain modulation in explaining the orientation tuning of astigmatic vision was greater in the chronic astigmatism group than in the induced astigmatism group, over an extended period (Figure 3B; cluster-based permutation test; p-value for chronic VS induced < 0.001, from 213 to 517 and p < 0.001 from 528 to 600 ms; p_chronic_ < 0.001, from 72 to 600 ms; p_induced_ < 0.001, from 49 to 247 ms). Optics-based prediction explained a substantial amount of the orientation tuning responses of the two astigmatism groups, and they were comparable with each other (Supplementary Figure 6A; cluster-based permutation test; p-value for chronic VS induced > 0.05 for whole duration; p_chronic_ < 0.001 from 100 to 531 ms; p_induced_ = 0.007, from 83 to 443 ms, p_induced_ = 0.027, from 485 to 600 ms). Therefore, the distorted retinal inputs from astigmatic vision were similarly represented in the EEG activity of both the induced and chronic astigmatism groups, while push-pull gain modulation estimated from the EEG activity was stronger in the chronic astigmatism group. Furthermore, in the induced astigmatism group, perceptual bias in their behavioral report correlated with optics-based predictions of the population tuning curve (Supplementary Figure 6B, right panel; p < 0.01 from 159 to 205 ms, and p < 0.05, from 327 to 342 ms), confirming that optics-based predictions accurately reflect perceptual fluctuations.

This gain modulation also had functional relevance to individual differences in perceptual decision-making. When we divided the participants in the chronic group into two based on the size of their average perceptual compensation (orientation bias reduction), participants with larger perceptual compensation exhibited greater gain modulation than those with smaller compensation (Figure 3C; cluster-based permutation test; p-value for low vs high < 0.001, from 126 to 484 ms). The levels of gain modulation and the extent of perceptual compensation were also significantly correlated with each other across participants (Supplementary Figure 6B, left panel; p < 0.05, from 132 to 293 and from 364 to 400 ms). On the other hand, this relationship did not hold in the induced astigmatism group (Supplementary Figure 7; cluster-based permutation test; p-value for low VS high > 0.05, for the whole duration). Additionally, fluctuations in perceptual compensation across different orientations (as shown in Figure 1C) were linked to changes in gain modulation. In the chronic astigmatism group, gain modulations across orientations exhibited a positive relationship with the reduction in behavioral bias for each participant (Supplementary Figure 6C, p < 0.001, from 131 to 423 ms). In contrast, no significant relationship was observed in the induced astigmatism group (Supplementary Figure 6D, p > 0.05 for all time points). These results demonstrate that the gain modulation observed in the posterior region of the brain is closely related to the restoration of orientation perception in the chronic astigmatism group.

One important point not to overlook is the presence of transient gain modulation in the induced astigmatism group (Figure 3B, right panel). Although this gain modulation was short-lived and showed no functional relevance to perceptual compensation (Supplementary Figure 6C, right panel, Figure 7, and Figure 9), it is noteworthy that such modulation can emerge even after relatively short-term exposure to astigmatic distortion. Approximately one hour of exposure was sufficient to induce compensatory neural gain modulation (see Figure 5A), but this effect did not persist over time (only briefly appeared between 49 and 247 ms from stimulus onset, Figure 3B). This largely overlaps with the time window (100–200 ms) during which the brain responds to contrast decreases caused by optical blur (Supplementary Figure 3), suggesting that the compensation observed in the induced astigmatism group was focused on contrast distortions. This suggests that early contrast compensation in the induced group must undergo a critical shift to the sustained orientation-specific compensation seen in the chronic group in order to influence perception.

### Inter-areal information transfer for neural compensation in astigmatic vision

To uncover the origin of the neural compensation in chronic astigmatism, we estimated the push-pull gain modulation across the whole brain by repeating GLM analysis on the neural orientation tuning responses estimated from every possible subset of EEG electrodes (Figure 4A). The gain modulation appeared first in the posterior electrodes, then appeared later in more anterior electrodes, as if it spreads gradually from posterior to anterior electrodes over time. To better understand the temporal dynamics of the inter-areal transmission of the gain modulation, we investigated the temporal updating of the gain modulation using a dynamical systems approach (Mante et al., 2013). Specifically, we focused on the transition matrices explaining the evolution of the push-pull gain modulation across time, where each element in the matrix represented the moment-to-moment transfer strength of gain from one electrode to another (see *Methods*, especially Equation 5). Thus, the transition matrix can reveal the transference of gain modulation in each electrode pair.

**Figure 4.**
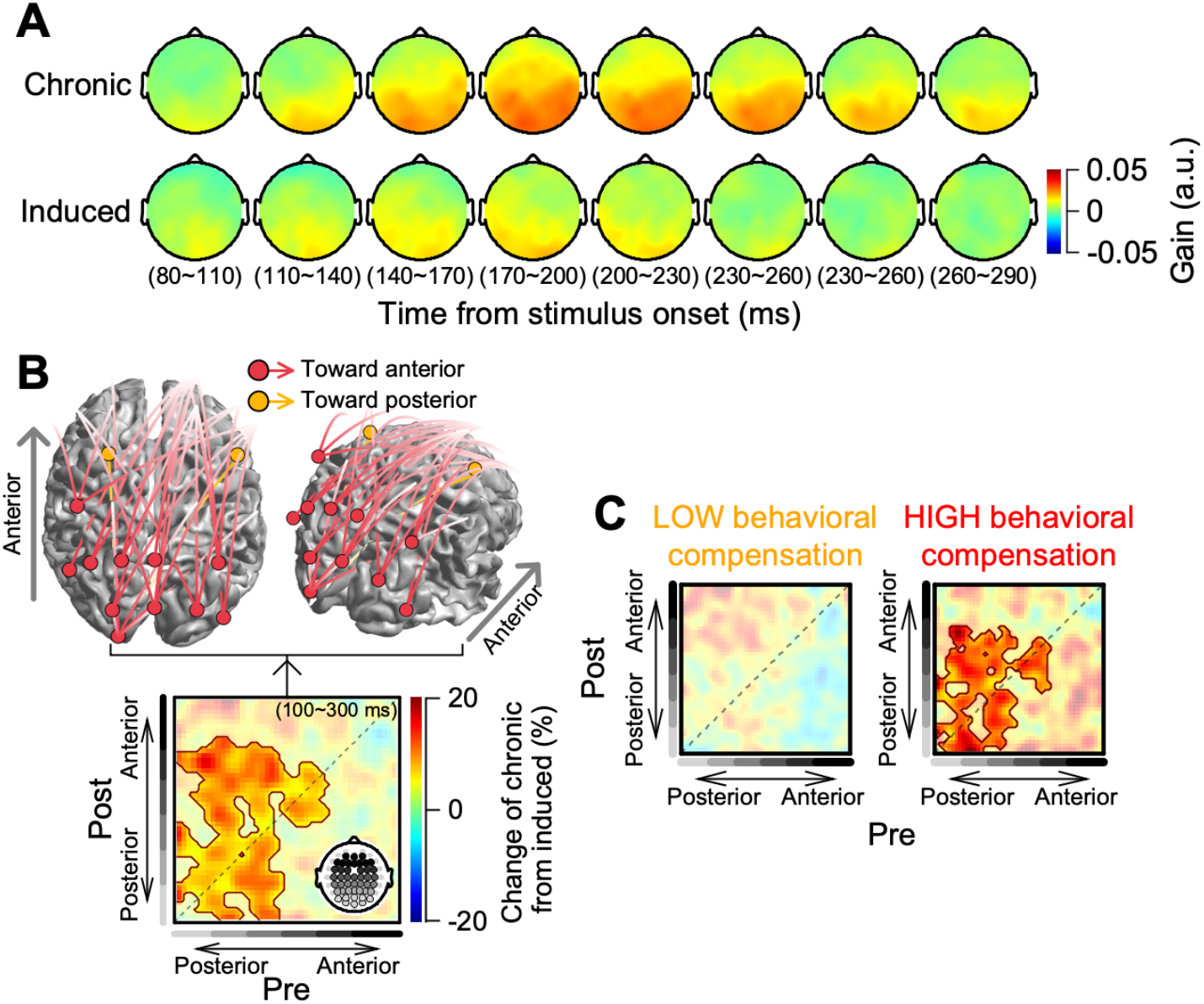
Transference of gain modulation in the chronic astigmatism group. **(A)** In the chronic astigmatism group, the searchlight approach unveiled a broad distribution of gain modulation originating from the posterior channels. **(B)** Bottom: Temporal transition of push-pull gain modulation across channels in the chronic astigmatism group compared to the induced astigmatism group. Each element represents the relative strength of gain transfer from an electrode in the corresponding column at a specific time point (*pre*-channel) to another electrode in the corresponding row at the subsequent time point (*post*-channel). Significant values are highlighted by contours around the red elements (p < 0.05). Values were averaged between 100 and 300 ms after stimulus onset. The brightness of the gray bars alongside each row and column of the matrix represents the spatial position of the channel along the anterior-posterior axis, as indicated by the inset topography (lighter gray: more posterior channels, darker gray: more anterior channels). Top: The transition matrix from the bottom subfigure was visualized by plotting pre-channels as circles on a hypothetical brain, with lines connecting them to their respective post-channels. Gain modulation transference in the posterior-to-anterior direction (upper triangle of the bottom matrix) is marked in red, while transference in the anterior-to-posterior direction (lower triangle of the bottom matrix) is marked in orange. Only transference exhibiting more than a 13% relative change is illustrated. **(C)** The strength of the push-pull gain modulation transition varies based on the degree of behavioral compensation.

Overall, the temporal gain transfer across recording electrodes was significantly greater than chance (Supplementary Figure 8A). To compare the transference of the gain modulation between the astigmatism groups, we calculated the percent change by taking the ratio of the transition matrices from the two groups. Notably, there was a more pronounced gain transfer from the posterior electrodes to other regions of the brain in the chronic astigmatism group compared to the induced astigmatism group (Figure 4B; cluster-based permutation test, ps < 0.001; average of 8.3% ± 1.2%, also refer to the upper triangle in Supplementary Figure 8A for absolute values of the transition matrix). The gain transfer from anterior to posterior regions was similar between the groups (as seen in the lower triangle of the transition matrix).

Furthermore the strength of the gain transfer in the chronic group has functional relationship with perceptual orientation compensation. When categorizing the chronic astigmatism participants by the degree of their behavioral compensation (Figure 3C), those with higher perceptual compensation exhibited a stronger posterior-to-anterior gain transfer than those of counterparts with lower behavioral compensation (Figure 4C; cluster-based permutation test, p = 0.005, relative gain transfer in comparison to the gain transfer seen in the induced astigmatism group). Such findings point to the role of gain modulation from the brain’s posterior region in mitigating biases in orientation perception.

As previously noted, gain modulation in the induced astigmatism group began to emerge after approximately one hour of exposure to the astigmatic condition. However, this effect did not persist throughout the trial (Figure 5A; cluster-based permutation test, p_1st half_ > 0.05 for the whole duration, p_2nd half_ = 0.002 from 95 to 243 ms), nor was it associated with perceptual orientation compensation (Supplementary Figure 9; cluster-based permutation test, ps > 0.05 for the whole duration). To understand the spatio-temporal structure of the neural gain modulation in the induced astigmatism group, we examined the changes in gain modulation transference by comparing the transition matrices estimated from the first and second halves of the experiment (percent change calculation). The transient gain modulation that emerged in the second half was accompanied by the anterior-to-posterior gain transferences—opposite in direction to the posterior-to-anterior transference observed in the chronic astigmatism group (Figure 5B; the lower triangle of the transition matrix; cluster-based permutation test, p = 0.002; refer to the lower triangle in Supplementary Figure 8B for absolute values of the transition matrix). In contrast, no significant differences in gain transference were observed in the chronic astigmatism group when comparing the first and second halves of the experiment (Supplementary Figure 8C; cluster-based permutation test, ps > 0.05).

**Figure 5.**
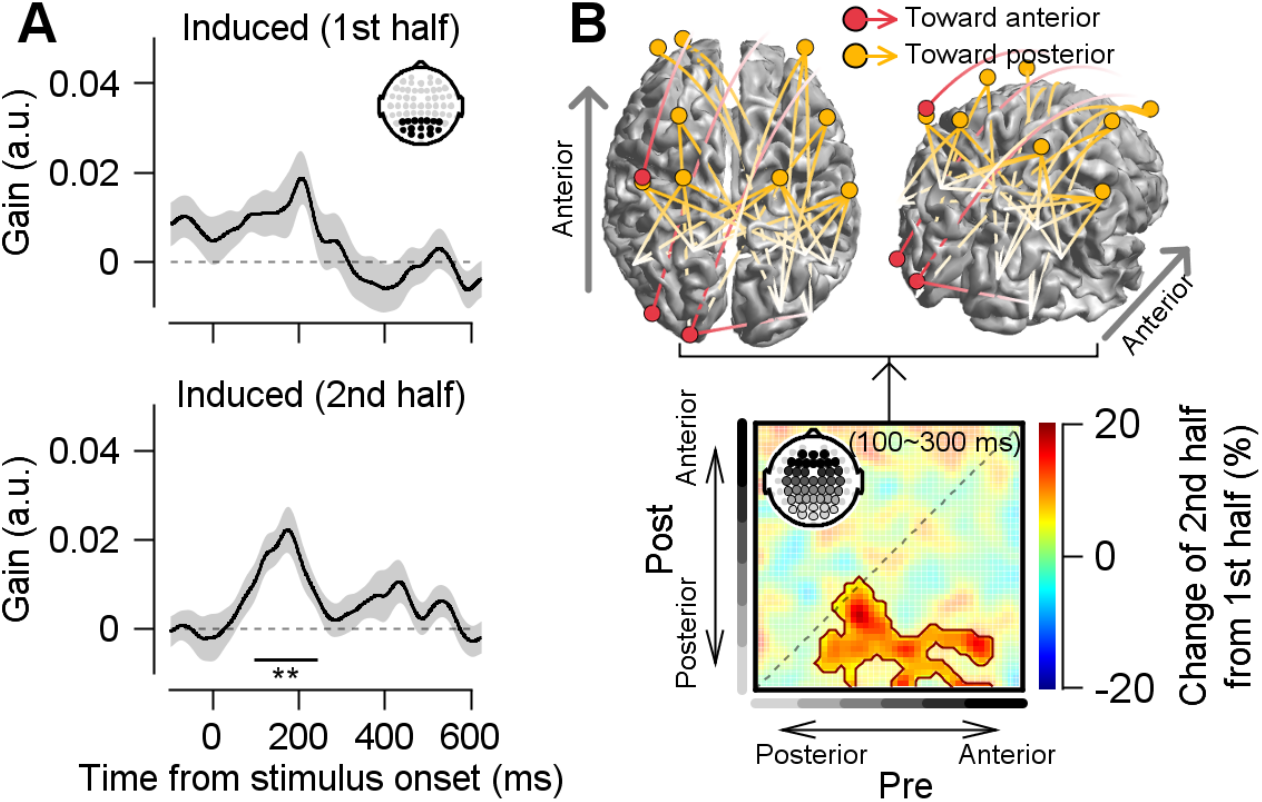
Gain modulation transference in the induced astigmatism group. **(A)** Gain modulation estimated from the model orientation tuning responses was computed separately for the first and second halves of the experiment (each lasting approximately one hour). The notations are identical to those in Figure 3. **(B)** Relative changes in the temporal transition of push-pull gain modulation across channels, comparing the first half with the second half of the experiment (bottom), along with its visualization (top). The notations are identical to those in Figure 4B.

The differences in inter-areal connectivity suggest that the posterior-to-anterior progression of gain modulation observed in the chronic astigmatism group emerges only after long-term exposure to astigmatic vision. Short-term exposure (approximately an hour in our experiment) induces transient neural gain modulation and the anterior-to-posterior gain transference, but without corresponding behavioral compensation. Chronic exposure, on the other hand, appears to drive plastic changes in posterior brain regions, optimizing orientation representation to more closely align with the physical world. Once this change is firmly established, the gain modulation that supports the restoration of orientation information may be systematically transmitted to other brain regions—ultimately influencing perceptual experience.

We note that the transference of gain modulation that we reported here is a relative measure. The anterior-to-posterior direction of gain transference in the induced astigmatism condition was relative to the first half of the induced condition, and the posterior-to-anterior direction of gain transference in the chronic astigmatism condition was relative to those in the induced condition. Although we cannot claim the absolute changes in gain modulation propagation, our results still provide insight into how this gain modulation spreads over the entire cortical surface as a function of how long we experience astigmatic vision.

## DISCUSSION

We investigated the neural mechanisms that optimize orientation perception in chronic astigmatic vision. We found that the stimulus orientation information represented in multivariate EEG activity was largely recovered despite the optical distortion caused by the chronic condition. The push-pull gain modulation systematically increased the neural discriminability of stimulus orientations near the astigmatic axis (the most blurring axis) and decreased the discriminability of the orthogonal orientations. Additionally, we demonstrated that the gain modulation for optimizing the distorted visual input was transferred from the posterior brain regions to the rest of the brain in the chronic condition, which was opposite from the transference direction in transiently induced astigmatism. Both the neural push-pull gain modulation and the strength of the gain modulation transfer in chronic astigmatism were correlated with perceptual compensation. This study shows the algorithmic principle by which the brain adjusts distorted sensory information based on the duration of exposure to the sensory deformation.

### Cortical neural mechanisms of gain modulation

Among the various mechanisms for achieving gain modulation(Bavelier et al., 2010; Fellous et al., 2003), the current study hypothesized that chronic astigmatic vision involves effective adjustments of single-neuron responses to orientations near the astigmatic axis—adjustments that cannot be solely explained by neurons’ responses to deformed retinal input. However, since we estimated orientation tuning responses from multivariate EEG activity based on neural discriminability across different stimulus orientations, the single-neuron mechanisms assumed in our model may not be the only contributors to the restoration of orientation tuning observed in EEG activity. In addition to the neural response amplitude adjustment assumed in our model (i.e., enhancing signal strength) (Murphy & Miller, 2009), improvements in neural orientation discriminability could also be achieved by suppressing background activity for orientations near the astigmatic axis (i.e., reducing neural noise) (Chance et al., 2002; Fellous et al., 2003).

We tested this alternative possibility by measuring the trial-by-trial variation of EEG activity and comparing it across different stimulus orientations (Supplementary Figure 5). We found that EEG activity variability did not significantly change under these conditions, which suggests that selective neural noise reduction (including intrinsic neural variability changes and interneuronal correlation changes (Nienborg et al., 2012)) might not be the underlying neural mechanism of the gain modulation estimated from EEG activity. Therefore, we believe that the amplification of single-neuron activities whose preferred orientations align with the astigmatic axis might be the tentative single-neuron level mechanism, as we suggested.

### Short-term and long-term plasticities in astigmatic vision

The underlying neural mechanisms of the push-pull gain modulation proposed in this study align with previous research on the effects of missing or weakened visual features (e.g., visual contrast deprivation) on perception. Prior studies have demonstrated a selective gain increase in neurons whose inputs were weakened due to environmental changes (Bao & Engel, 2012; Haak et al., 2014; Zhang et al., 2009). Specifically, when participants were exposed to environments where a specific orientation feature was artificially removed, perceptual compensation for the weakened orientation was attributed to increased selective responsiveness of visual cortical neurons to the weakened input (Zhang et al., 2009). In our optics-based model, the push-pull gain modulation implemented this selective responsiveness adjustment by enhancing the response sensitivity of neurons whose preferred orientations aligned with the astigmatic axis—where optical blur is maximized. Furthermore, we validated our model by comparing its predictions with the orientation tuning functions estimated from multivariate EEG activity patterns.

How does this modification of visual cortical neural activity occur? This is a difficult question to answer, given that we do not have direct, cellular-level measures of neural compensation. However, we can infer some aspects of this process from the neural compensation observed in the induced astigmatic vision condition. In the induced condition, we did not expect any neural compensation to occur because the study was not initially designed to investigate short-term plasticity. Although we did not control the exposure duration, prolonged exposure to the induced condition (about an hour of intermittent exposure) resulted in neural plasticity, with transient orientation compensation emerging (Figure 3B, right panel).

While the gain modulation observed in the chronic condition was maintained for an extended duration (Figure 3B, left panel), the gain modulation in the induced condition appeared briefly and did not impact perceptual compensation (Supplementary Figure 9). This difference might suggest the existence of separate neural mechanisms for short-term and long-term adaptation (Bao & Engel, 2012; Haak et al., 2014). The transient gain modulation might be shared across the two adaptation processes, but maintaining it for an extended period could be crucial for perceptual adjustments.

Therefore, the extended duration of neural gain modulation might serve as a neural signature of the network state, marking the transition from flexible short-term plasticity to stabilized long-term plasticity with established roles in perception (Lazzouni & Lepore, 2014; Liu et al., 2007). Once the neural network stabilizes, neural gain modulation in the visual system could propagate to other brain regions (Figure 4), enabling perceptual compensation to become automatic after prolonged exposure to visual impairments (Barbot, Park, et al., 2021; Mitchell et al., 1973; Son et al., 2022).

In contrast, during short-term adaptation in the induced condition, transient gain modulation might result from active cognitive control originating in frontal cortical brain regions (Figure 5B). We note that exposing participants to the induced astigmatic vision condition for a longer duration and in a more controlled manner might allow observation of the actual transition of gain propagation from the anterior-to-posterior direction to the posterior-to-anterior direction, as well as the emergence of extended-period gain modulation during each trial. This could be an important line of research in the future, to understand how fast and slow neural processes interact to contribute to perceptual compensation.

### Cognitive processes and top-down modulation

The compensatory gain modulation we observed shares similarities with the neural mechanisms underlying perceptual learning (Bavelier et al., 2010). Specifically, push-pull gain modulation has been demonstrated in perceptual learning studies (Li, 2016; Watanabe & Sasaki, 2015; Yan et al., 2014), where plastic changes occur in visual cortical neural activity and local circuits (Jehee et al., 2012; Karni & Sagi, 1991; Sanayei et al., 2018; Schoups et al., 2001; Yang & Maunsell, 2004). However, in perceptual learning, positive feedback enhances sensitivity to frequent stimuli, while chronic astigmatism reduces sensitivity to frequent orientations due to years of indirect negative feedback from the mismatch between perception and reality. This mismatch may arise from processes such as reorganizing the selectivity of neural populations for frequent stimuli to reinforce weaker ones (Ajina et al., 2021; Baker et al., 2005; Barbot, Das, et al., 2021) or neural adaptation to those frequent stimuli (Son et al., 2022; Webster et al., 2002). This suggests that neural compensation for astigmatism may result from extensive daily perceptual learning.

Another critical cognitive process likely triggering the compensation process is top-down modulation. Upon noticing the mismatch between reality and perception under astigmatism, feedback from frontal cortical areas may emerge along the hierarchy of the visual system, similar to that during perceptual learning (Byers & Serences, 2014; Dosher & Lu, 2017; Jing et al., 2021; Moldakarimov et al., 2014; Yang & Maunsell, 2004) and top-down attention (Armstrong et al., 2006; Cavanaugh et al., 2019; Moore & Armstrong, 2003). This top-down modulation may be the neural origin of the short-term plasticity observed in the induced astigmatism condition — the propagation of the compensatory gain modulation from anterior to posterior regions in the induced astigmatism (Figure 5). The compensatory effect was highly time-selective, lasting only 100–200 ms after stimulus presentation. Notably, this period is primarily associated with contrast-related neural responses in individuals with visual impairments (Regan, 1973; Sokol, 1983; Son et al., 2021). This may suggest that extra-sensory cortices detected environmental irregularities — specifically, weakened contrast for a particular visual feature — and triggered compensatory feedback to the visual cortex.

### Practical implications

Lastly, our findings highlight the crucial role of neural plasticity in addressing visual impairments in practical applications. When complete optical correction for visual impairments, including astigmatism, is performed without considering extra-retinal factors—whether through eyeglasses, surgery, or other interventions—it can lead to performance decline (Barbot, Park, et al., 2021; Ng et al., 2021), perceptual shifts toward near- or far-sightedness in clinical practice (Benavente-Pérez et al., 2014; McLean & Wallman, 2003), and even unexpected astigmatic symptoms with a reversed blurring axis (Mitchell et al., 1973; Son et al., 2022). This raises the challenge of determining the appropriate degree of optical correction (Villegas et al., 2014).

Neural plasticity following surgical interventions such as LASIK (laser-assisted in situ keratomileusis) or LASEK (laser-assisted sub-epithelial keratectomy) serves as a compelling example, as these procedures aim for complete optical correction but may result in suboptimal perceptual experiences (Kim et al., 2019). Based on our findings, we emphasize that plastic changes in the brain must always be considered in the context of visual correction. This neural constraint also has significant implications for the development of peripheral prostheses; even state-of-the-art retinal prostheses can only restore missing visual input, which does not inherently guarantee accurate perceptual experiences (Beyeler et al., 2017; Castaldi et al., 2020). The present study underscores the need for a comprehensive understanding of neural compensation mechanisms to optimize optical correction strategies.

## METHODS

### Two groups of participants: chronic and induced astigmatism

Data were collected from 42 participants (27 males and 15 females; mean age, 22.7±3.4). The refractive error of the naked dominant eye was measured using an auto refractometer (AR; Huvitz, HRK-7000, Republic of Korea). The dominant eye was identified based on monocular preference in binocular viewing (Purves & White, 1994). Nineteen participants were classified into the induced astigmatism group (normal-vision) since their spherical and cylindrical refractive errors were both within the 0.75 diopter range from the emmetropic state (mean refractive errors, spherical, 0.20±0.27 diopter, cylindrical, 0.28±0.30 diopter). The remaining refractive errors would be negligibly small (see Table 20-1 in the reference (Benjamin & Borish, 2006)), and fully corrected before the experiment. Twenty-three participants were classified into the chronic astigmatism group as their cylindrical refractive errors were +1.00 diopter or worse (mean cylindrical refractive errors, 2.36±1.11 diopter; see Table 1 for the detailed information of each participant). The Institutional Review Board of Sungkyunkwan University approved the study (IRB 2018-05-003), and all studies followed relevant guidelines. All participants provided written informed consent and were naive to the purpose of the experiment.

### Preparation for astigmatic vision condition

Comparable myopic regular astigmatism was applied to all participants; the light rays in a certain meridian focused before the retina as shown in Figure 1A. In both groups, we fully corrected the spherical refractive errors of the test eye so that the near-sightedness or far-sightedness would not interfere their vision. In the chronic astigmatism group, their innate cylindrical refractive errors of the test eye were left uncorrected so their visual perception would be dominated by their own astigmatism (Supplementary Figure 2A, upper left panel). In the induced astigmatism group, we fully corrected their innate astigmatism and placed an additional +2.00 diopter cylindrical lens to make them have a similar experience to the chronic astigmatism group (Supplementary Figure 2A, lower left panel). In the induced astigmatism group, if participants had mild cylindrical refractive errors (≥ 0.25 diopters), the cylindrical lens axis was matched to the angle indicated by the auto-refractometer. For those with no such errors, the axis was set horizontally, based on the typical pattern in the chronic astigmatism group, which predominantly exhibited ‘with-the-rule astigmatism,’ characterized by strongest refraction along the vertical meridian. Additionally, to make the light rays penetrate the lens parallel at the viewing distance, we added a +1.50 spherical lens in front of all test eyes. We placed a +10.00 diopter spherical lens to the opposite eye of the tested one, to prevent undesired intervention. We also obtained data from each participant’s emmetropic states of the identical eye by fully correcting both spherical and cylindrical refractive errors (Supplementary Figure 2A, right panels). The participants were instructed to keep both sides of their eyes open and place their chin on a chin-rest and their forehead against a forehead-rest. This setup helped to reduce potential confounding factors by maintaining a consistent viewing distance and limiting head movements such as turning or tilting.

### Task design

Participants were instructed to report the perceived mean orientation of a tilted Gabor array, which was presented for 153 ms (Figure 1B). The array consisted of non-overlapping 20 Gabor stimuli within an invisible 2.5° radius circle located at the center of the screen. Each Gabor with a 0.25° radius had a spatial frequency of 2 cycles/° and randomly selected polarity of either 0° or 180° with a Michelson luminance ratio of 60%. All Gabor stimuli were tilted identically from the measured astigmatic axis, and the tilt was randomly selected from eight orientations for each trial (0°, ±22.5°, ±45°, ±67.5°, or 90°). After 494 ms of the post-stimulus blank screen, the participants were asked to rotate a white bar to report the perceived orientation of the array by moving the mouse and saved their response by clicking the mouse button. The participants pressed the space bar to initiate the next trial. A white fixation point with a radius of 0.1° appeared from the moment the space bar was pressed until the white response bar (with a radius of 5° and a Michelson luminance ratio of 100%) turned on, and the participants were asked to fixate on it. We used small, low-contrast Gabor patches as target stimuli for orientation perception to maximize the effect of astigmatic distortion. This design required participants to rely on the individual luminance edges of each Gabor to determine orientation. In contrast, the large, high-contrast white response bar exhibited strong resistance to astigmatic distortion, as its overall orientation information remained intact despite systematic distortions in the luminance edges (Supplementary Figure 1C).

Each participant performed eight blocks of a total of 1280 trials. Half of the block was obtained under the astigmatic vision condition (as Supplementary Figure 2A left panels; for details, see Preparation for astigmatic vision condition section), and the other half after all refractive errors were fully corrected (emmetropic state). We alternated block types without informing the participants. Each block immediately began after they put on the glasses corresponding to that block’s condition. Participants wore the specific lens for an average of 10 to 15 minutes in each block. This was to minimize any uncontrolled improvements in visual acuity that could result from adaptation to the optical settings (Vinas et al., 2012; Yehezkel et al., 2010).

All visual stimuli were presented on a gamma-corrected, 20-inch CRT monitor with a spatial resolution of 800 × 600 pixels and a vertical refresh rate of 85 Hz (Hewlett Packard p1230; maximum and minimum luminance of 106.0 cd/m^2^ and 0.8 cd/m^2,^ and gray background luminance 55.8 cd/m^2^) in the dark room. All the stimulus presentations were controlled by a custom program written in MATLAB (Mathworks Inc.) using the Psychophysics Toolbox (Brainard, 1997; Pelli, 1997).

### Response bias and compensation

Reported perceptual orientation biases were quantified in the following way. The response bias was first obtained by subtracting the stimulus orientation from the reported orientation. Then, we reversed the sign when the stimulus orientations were between 0° and 90°. Therefore, a positive astigmatic bias means the reported orientation is biased away from the astigmatic axis, and a negative bias indicates the reported orientation is biased toward the astigmatic axis. The astigmatic biases were averaged according to the deviation from the astigmatic axis (Figure 1C). We used the circular statistics toolbox for MATLAB (Berens, 2009) to calculate all the response biases.

To further derive the behavioral adjustments that were higher or lower than the bias predicted from the optical distortion (see Optics-based model prediction section for more details), we obtained the difference between the measured behavioral bias and the bias predicted from the optics-based model. A positive adjustment means a smaller perceptual bias than that from the model prediction, and a negative adjustment means a larger bias than that from the model prediction. The overall amount of behavioral compensation for each participant is shown in Figure 3C (left panel, chronic astigmatism group) and Supplementary Figure 7 (left panel, induced astigmatism group).

### Optics-based model prediction

We built a model predicting the population orientation tuning curve in the brain and the behavioral responses purely driven by the measured refractive errors. First, we estimated the optical blur using the relative length of the retinal image in the astigmatic vision state compared to the emmetropic vision state (λ), given each eye’s optical property of cylindrical refractive error.

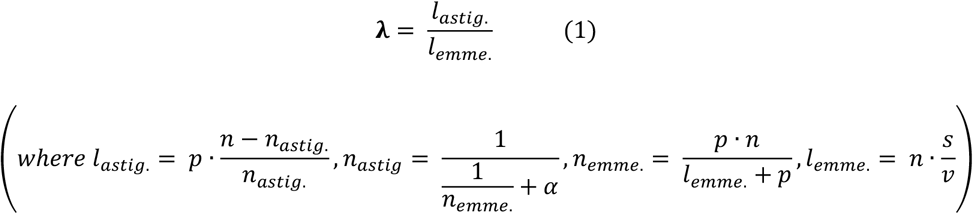

*l*_*astig*._ and *l*_*emme*._ denote the length of the retinal images in astigmatic and emmetropic eyes, respectively (see Supplementary Figure 2B for a graphical illustration). *n, n*_*astig*_., and *n*_*emme*._ represent the nodal distance and the distances between the nodal point and the focal plane in astigmatic or emmetropic eyes, repectively. The nodal distance (*n*) and pupil size (*p*) were assumed to match typical adult eye dimensions (*n*=24mm, *p*=3.5mm). The variable *α* refers to the eye’s cylindrical refractive error, measured in diopter using an auto-refractometer. *s* and *ν* represent the actual pixel size of the diplay (*≈*0.50 mm) and viewing distance (60 cm), respectively. Equation 1 was derived using the *Vertical Angles Theorem* and the *Proportionality of Similar Triangles*, by solving the geometric relationship between screen pixel size, viewing distance, nodal distance, and changes in nodal distance due to additional astigmatic refractive power.

In the second step, we generated the predicted retinal image (*I*) of the visual stimulus based on the estimated degree of the optical deformation. This was done by convolving the Gabor stimulus image with an oval-shaped kernel (Figure 2A, left panel). The horizontal diameter of the kernel was fixed at 1 pixel, while the vertical diameter was scaled according to the image length ratio calculated in the first step (i.e., *l*_*astig*._*/l*_*emme*._). This procedure simulates astigmatic optics by introducing elliptical spreading of light rays—mimicking the blur distortion observed in astigmatic vision (compare the shape of a light ray before and after the lens in Figure 1A).

To assess potential confounding factors and validate Equation 1 of the optics-based model, we also utilized the open-source optics simulation program ISETBIO (Cottaris et al., 2019) (Supplementary Figure 1B) to simulate astigmatism-induced retinal image distortion. The retinal image of a single Gabor graiting stimulus was generated using the following parameters: a 0.5-degree field of view, wavelengths ranging from 400 to 700 nm in 10 nm increments, a pupil size of 3.5 mm, and each participant’s individual cylindrical refractive error.

As the third step, we estimated the population tuning responses 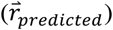 based on the assumption that the tuning function reflects the collective response of hypothetical neuronal units with varying orientation preferences. The predicted population response of hypothetical neurons with a preferred orientation *θ* to a given visual stimulus is defined as follows:

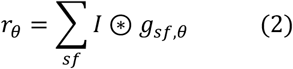

Here, the retinal image (*I*, estimated from the second step) was convolved (⊛) with kernel filters (*g*_*sf*_,_θ_) tuned to a preferred orientation *θ* and three spatial frequencies (*sf:* 1, 2, and 4 cycles per degree, see Figure 2A, left panel for sample illustrations of these filters). The convolved data for a given preferred orientation were then integrated across all three spatial frequencies. Multiple spatial frequency filters were used to account for changes in the stimulus’s effective spatial frequency in the retinal image, which can occur due to astigmatism-induced blurring in the second step. Next, this procedure (as described in Equation 2) was repeated for each preferred orientation of the hypothetical neuronal units, resulting in the predicted population tuning response vector 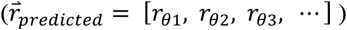. This approach closely resembles the computation of population orientation tuning curves derived from simple cell activities in the primary visual cortex (Bloem & Ling, 2019; Hubel & Wiesel, 1959; Jones & Palmer, 1987; Lee, 1996; Portilla & Simoncelli, 2000).

We adjusted the predicted population tuning curve to place the responses of the model units, whose preferred orientation matched with the stimulus orientation, at the center of the tuning curve (Figure 2A, middle right panel). This ensured that responses of model units, whose preferred orientations deviated from the stimulus orientation, were located at the flank of the population tuning curve (a process we refer to as “zero-centering” based on the actual stimulus orientation). For the population tuning responses to the stimulus orientations between 0° and 90°, we used the mirror image of the tuning curve to ensure that the model response to the orientation of the astigmatic axis consistently appeared on the right side of the population tuning curve (Figure 2A, middle right panel). Finally, we calculated the weighted circular mean of the zero-centered tuning function, using the amplitude at each orientation as the weight. This circular mean represents the center of mass of the orientation tuning function. The angle difference between this center and the veridical stimulus orientation was taken as the model’s predicted perceptual bias (dashed lines Figure 1C and right panel of Figure 2A). All calculations were performed using the circular statistics toolbox for MATLAB (Berens, 2009).

### EEG data acquisition and preprocessing

During the orientation perception task, we collected 64-channel EEG activity (actiCAP, Brain Products, GmbH) using the amplifier (BrainAmp, Brain Products, GmbH) with a sampling rate of 5 kHz. We maintained the impedances of all the electrodes below 15 kΩ during the recordings. The collected EEG data were preprocessed in the following order. First, they were down-sampled to 1 kHz and then high-pass filtered with a 2^nd^ order Butterworth filter at a cut-off frequency of 0.1 Hz. We removed noisy channels using Artifact Subspace Reconstruction (ASR) (Mullen et al., 2015), and then re-referenced the EEG data using the average as the reference (Bigdely-Shamlo et al., 2015). We further removed the remaining line noise of 60 Hz and its harmonics with the *cleanline* EEG Lab plugin. Finally, artifactual components identified by independent component analysis (ICA) (Makeig et al., 1995) were rejected automatically by the ADJUST EEG Lab plugin(Mognon et al., 2011).

Event-related potentials (ERPs) were obtained by averaging EEG activity from the Oz, Pz, Cz, and Fz electrodes separately for each participant under astigmatic and emmetropic vision conditions (Supplementary Figure 3A) and for each stimulus orientation presented under the astigmatic vision condition (Supplementary Figure 3B).

The standard deviation of EEG activity from posterior electrodes was measured for each stimulus orientation and realigned so that the values corresponding to the stimulus orientation of the astigmatic axis were centered (Supplementary Figure 5).

### Orientation tuning response

We derived the orientation tuning response (Figure 2B-D) from the EEG activity recorded at the posterior electrodes. In each time point, we calculated the Mahalanobis distances between the multivariate EEG activity pattern for a specific orientation stimulus on a given trial and the average activity patterns from eight different stimulus orientations. This produced an 8-by-1 distance vector. (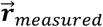 in Equation 3 below) (Myers et al., 2015; Wolff et al., 2015, 2017). We reorganized the distance vector to position the distance value for the given stimulus orientation at the center.

Consequently, distance values for the other orientations radiated out from this central point, a process we refer to as “zero-centering”. After subtracting the mean, the sign of the distance vector was reversed to make higher EEG activity pattern similarities have higher values. We analyzed the orientation tuning responses specifically for oblique orientations (±22.5°, ±45°, and ±67.5°) to assess the distortion of tuning functions under both astigmatic (Figure 2D) and emmetropic vision conditions (Supplementary Figure 4) across the astigmatism groups. It is important to note that we used the mirror image of the tuning response when the stimulus orientation was +22.5°, +45°, or +67.5°. This ensured the astigmatic axis consistently appeared on the same side of the tuning curve, which is a similar approach to what we took in calculating the model prediction of behavioral biases. This process was conducted separately for each participant’s data from astigmatic and emmetropic vision conditions.

We evaluated the bias in the orientation tuning estimated from the EEG activity pattern (Figure 2D and Supplementary Figure 4) by converting the tuning curve into a probability density function and estimating the skewness of the function (Olofsson, 2012). We also estimated the amplitude of the tuning response by convolving it with a cosine kernel raised at 15^th^ power (Myers et al., 2015).

When the orientation tuning function is estimated using this method, its center will exhibit the maximum value, as the distance between neural response patterns for the same stimulus is minimized. However, when retinal distortion occurs—particularly in the induced astigmatism condition—the estimated orientation tuning function becomes skewed, even though the peak remains centered, as shown in a previous study (Son et al., 2021). In some figures, we applied the Von Mises function to the average orientation tuning responses to enhance readability and visual clarity, which results in a shifted representation of the tuning function. Importantly, this fitting was used for visualization purposes only. All quantitative analyses—including accuracy and skewness estimation (Figure 2D), as well as the gain modulation model estimation—were based on the raw average orientation tuning responses.

We estimated orientation tuning responses using Mahalanobis distance within each astigmatism condition. One might consider an alternative approach—estimating orientation tuning in the astigmatic condition by calculating the neural distances between responses in the astigmatic vision condition and those in the emmetropic condition. This method could, in theory, provide a more direct measure of how much the orientation tuning response deviates from that of normal vision.

However, in multivariate EEG analysis, changes in experimental conditions can introduce unexpected shifts in the overall neural representational space. As a result, the neural orientation spaces across conditions may be misaligned or offset, making direct comparisons problematic. In such cases, proper scaling or normalization of the neural responses is necessary, as has been used in a previous study (Son et al., 2021). As an alternative, we estimated neural orientation tuning functions independently within each vision condition. This approach serves as an implicit normalization, enabling meaningful comparison of orientation tuning across astigmatism conditions without assuming that the representational spaces are directly aligned.

### Isolating extra-retinal information from orientation tuning response

We built a generalized linear regression model (GLM) whose measured orientation tuning response was assumed to be a summation of a component directly receives retinal inputs that are distorted by the astigmatic vision and the other component derived from an extra-retinal process that modulates the gain of the first component. The model is formalized as follows:

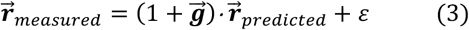

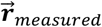 is an 8-by-1 vector representing the measured orientation tuning response, extracted from multivariate EEG activities and converted into a probability density function (see *orientation tuning response* section for details). 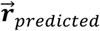 is an 8-by-1 vector of predicted orientation tuning response derived from the optics-based model for a given set of stimulus orientations. These predictions incorporate participant-specific retinal input based on individual optical properties and are converted into population orientation tuning functions (Figure 2A, left panel; see *Optics-based model prediction* for more detail). Note that 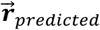 may vary across participants due to individual differences in optical characteristics. 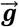 is an 8-by-1 vector representing the gain function, defined as the values of a cosine function across the eight stimulus orientations (Figure 3A, upper right panel). The dot (·) denotes element-wise multiplication, and *ε* represents the error term.

We modified Equation 3 to estimate the contributions of the distorted retinal input and extra-retinal gain modulation on the measured orientation tuning responses.

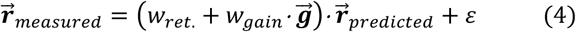

where *w*_*ret*._ is a scalar value that determines the weight of the retinal inputs and *w*_*gain*_ is a scalar value that adjusts the contribution of the gain modulation. We utilized a two-step approach to fit the three free parameters of the GLM model. Initially, we fitted *w*_*red*._ and *ε*, focusing exclusively on the theoretical predictions of the optics-based model. Subsequently, *w*_*gain*_ was fitted to the residuals from this initial model to capture additional variance attributable to neural gain modulation. This two-step strategy was adopted to prevent the model from overemphasizing *w*_*gain*_. Although we also tested a simultaneous fitting of all free parameters at once, the results were largely consistent with those obtained using the two-step approach (Supplementary Figure 6E).

We estimated the GLM by fitting it to the orientation tuning responses extracted from the posterior electrodes (Figure 3, Figure 5A, and Supplementary Figure 6A). Additionally, we estimated the GLM from subsets of electrodes (Figure 4A) using a localized analysis in which electrodes within 10% of the head radius surrounding each individual electrode were included in the orientation tuning estimation—an approach conceptually similar to the searchlight method (Hajonides et al., 2021; Kriegeskorte et al., 2006). Orientation tuning responses were individually estimated for 45 electrodes, excluding those located on the neck, around the ears, or directly above eyebrows.

### Behavioral relevance of gain modulation

We averaged neural gain modulation and behavioral compensation values—estimated for each stimulus orientation at each time point—across trials. Using these averages, we performed a simple linear regression with eight data pairs, corresponding to the eight stimulus orientations. In the regression, the neural gain modulation was plotted on the x-axis and the behavioral compensation on the y-axis. A positive regression slope indicates that stronger gain modulation in the tuning function— presumably driven by the extra-retinal process—is associated with greater behavioral compensation. Finally, we averaged the regression slopes across participants to evaluate the consistency and validity of this relationship within indivisuals (Supplementary Figure 6C-D).

In addition, we tested if the variation of the gain modulation across participants is correlated with the variation in behavioral compensation. We divided the participants in the chronic astigmatism group into the two according to the sizes of the behavioral compensation (median-split): a group with high behavioral compensation and the other with low compensation (Figure 3C, left panel). Then, we compared the two groups’ neural gain modulation observed in the orientation tuning responses (Figure 3C, right two panels). Additionally, the Spearman correlation coefficient between the amount of behavioral compensation and that of gain modulation at each time point was computed across participants (Supplementary Figure 6B, left panel). The significance of the correlation coefficient was assessed by comparing the observed correlation with a null distribution obtained through permutation analysis. In this analysis, the participant-wise mapping between neural gain modulation and behavioral compensation was disrupted by randomly shuffling participant labels 1000 times, generating a distribution of correlation coefficients under the null hypothesis. Lastly, we calculated the Spearman correlation between *w*_*ret*._ (the weight applied to the retinal input, estimated at each time point) in Equation 4 with perceptual biases of participants in the induced astigmatism group, to test if the optics-based model accurately predicted the perceptual fluctuation (Supplementary Figure 6B, right panel).

### Inter-regional connectivity within gain modulation dynamics

We analyzed the temporal transitions of gain modulation across electrodes to test if there are directional spatial spreading of neural gain modulation in the brain. For this purpose, we first defined a dynamical system of gain modulation using Equation 5 below.

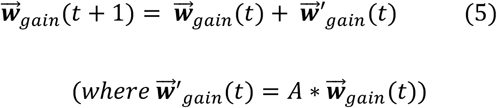

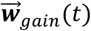 is a 45-by-1 vector consisting of weight values for the gain function (*w*_*gain*_ for each electrode was estimated from Equation 4) of all 45 electrodes at the time point *t*. The change of gain modulation from the time point *t* to 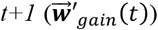 was the result of matrix product (*) between 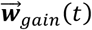 and the transition matrix A. An element of A at the i^th^ row and j^th^ column, *a*_*ij*_, represents the weight for the transition from the j^th^ electrode to the i^th^ electrode. In each time point, we estimated the temporal transition matrix by modifying Equation 5 into,

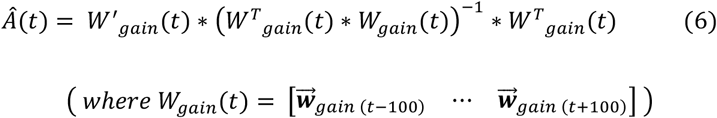

To estimate the transition matrix, we employed a five-fold cross-validation approach. Trials were randomly divided into five groups, with 80% of the trials used to estimate the transition matrix and the remaining 20% used to generate *W*_*gain*_ (the weights for gain modulation) based on the estimated matrix. We then assessed the reliability of this estimation by calculating the cosine similarity between the *W*_*gain*_ generated from the estimated transition matrix and the actual gain modulation dynamics from which the matrix was derived. The two time series showed a strong correspondence: the average cosine similarity over the 100–300 ms post-stimulus time window was 0.052 ± 0.006 (*t*_41_ = 8.897, *p* = 4.036e-^11^). This five-fold cross-validation procedure was repeated 10 times, and the resulting transition matrices were averaged across iterations. Finally, we took the absolute values of the averaged transition matrix to construct the connectivity matrix (Supplementary Figure 8A and B).

To test the significance of the connectivity matrix, we performed a permutation test. A null distribution was generated by repeating the transition matrix estimation procedure described above 500 times, each time using data in which the order of the gain function weights was randomly shuffled in both time and space (i.e. shuffled columns and rows of *W*_*gain*_*(t)*). This allowed us to evaluate whether the observed connectivity values significantly exceeded those expected by chance. We then computed *z*-scores for each element of the transition matrix by subtracting the mean of the null distribution and dividing by its standard deviation. The statistical significance of individual elements was evaluated by testing whether each *z*-scored element exceeded a threshold of 2 (i.e., > 2 standard deviations) using a cluster-based permutation test (Maris & Oostenveld, 2007).

Finally, we probed the relative change of the transition matrix in the chronic astigmatism group to that in the induced astigmatism group by calculating the ratio between the two (Figure 4B). We also assessed changes in the transition matrix within each astigmatism group by comparing the first and second halves of the trials, again using the ratio between the two (Figure 5B and Supplementary Figure 8C). Additionally, to evaluate the relevance of the transition matrix in relation to perceptual compensation, we divided participants in the chronic astigmatism group into two subgroups based on the magnitude of their perceptual compensation. We then compared the transition matrices between these two subgroups (Figure 4C).

## Author contributions

Conceptualization: SS, HK, and JL

Methodology: SS, HK, HRK, and WMS

Investigation: SS and HK

Formal analysis: SS

Writing—original draft: SS and JL

Writing—review & editing: SS, HK, HRK, WMS, and JL

## Data and materials availability

The datasets generated and/or analyzed during the current study are available at https://github.com/SangkyuSon/astigEEG. Correspondence and requests for materials should be addressed to SS and JL.

## SUPPLEMENTARY FIGURES

**Supplementary Figure 1.**
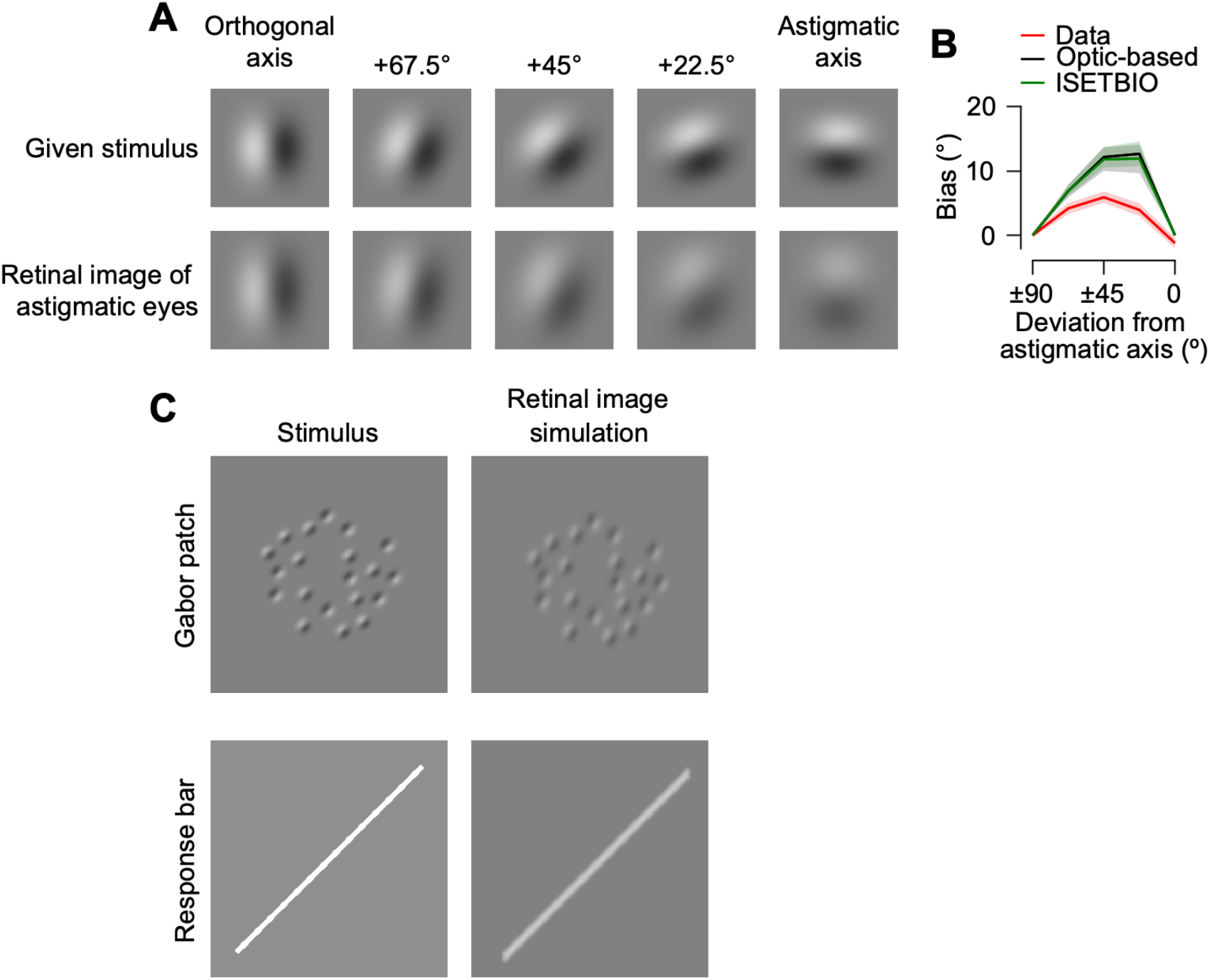
A simulated retinal image of astigmatic eyes. **(A)** Axis-specific optical blur was simulated by convolving the given stimulus image with an oval-shaped kernel. The degree of ellipticity was determined by each participatn’s cylindrical refractive error (see Equation 1). **(B)** Two models, the optics-based model used in the main text (black) and the open-source simulation model ISETBIO (green), predicted perceptual biases in the chronic group. Shaded regions represent ±1 SEM. **(C)** Simulated retinal images of the Gabor patch and response bar. The orientation of the Gabor patch depended on the short edges of individual stimuli, whereas the response bar’s orientation remained comparatively clear due to its higher contrast and global structure.

**Supplementary Figure 2.**
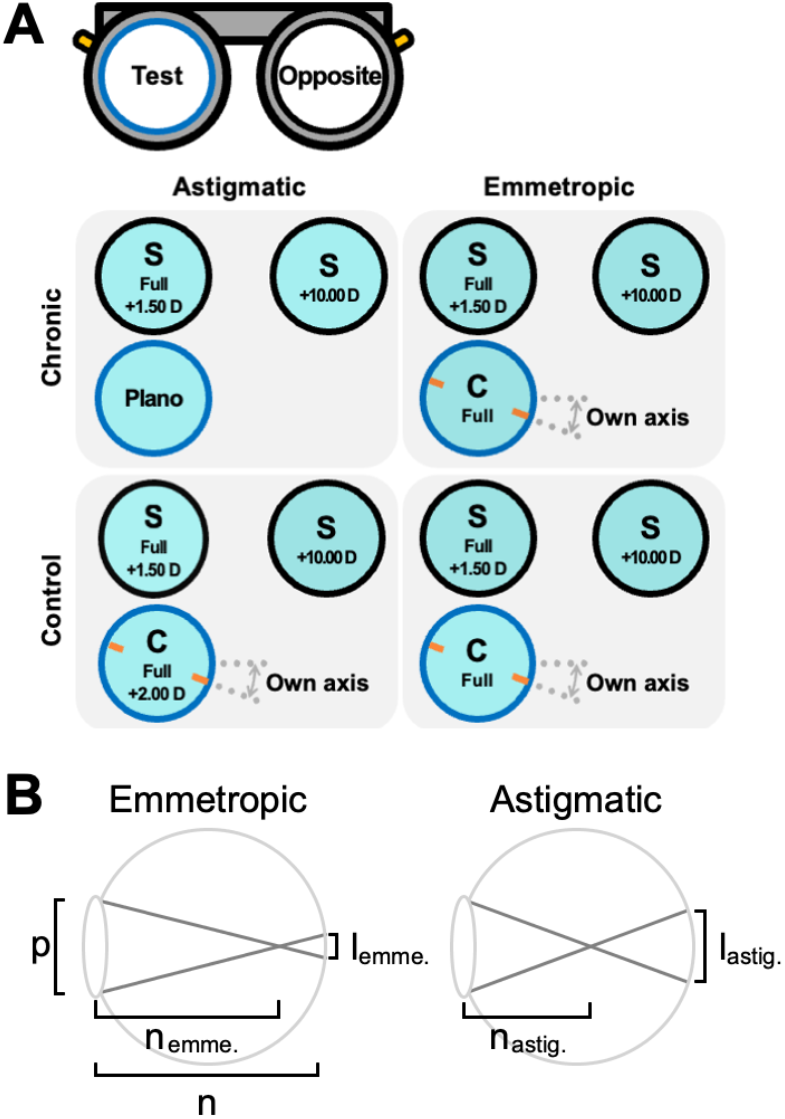
Experimental setting and optics-based model. **(A)** Each panel displays the trial lens composition layout in the chronic (upper row) or control (lower row) astigmatism groups under the astigmatic (left column) or emmetropic vision conditions (right column). The notations S, C, full, and D indicate a spherical lens, cylindrical lens, full correction, and diopter. Lenses with critical manipulations for each condition are highlighted with a blue line. **(B)** Illustration of how the retinal image was optically blurred in the optics-based model. *p, n, n*_*emme*_, *n*_*astig*._, *l*_*emme*._, and *l*_*astig*._ indicates pupil size, nodal distance, nodal point to focus in emmetropic vision, nodal point to focus in astigmatic vision, length of the retinal image in emmetropic vision, and length in astigmatic vision. In the astigmatic vision, since the light ray refracts more, it has a longer length of the retinal image (i.e., *l*_*astig*._ > *l*_*emme*._).

**Supplementary Figure 3.**
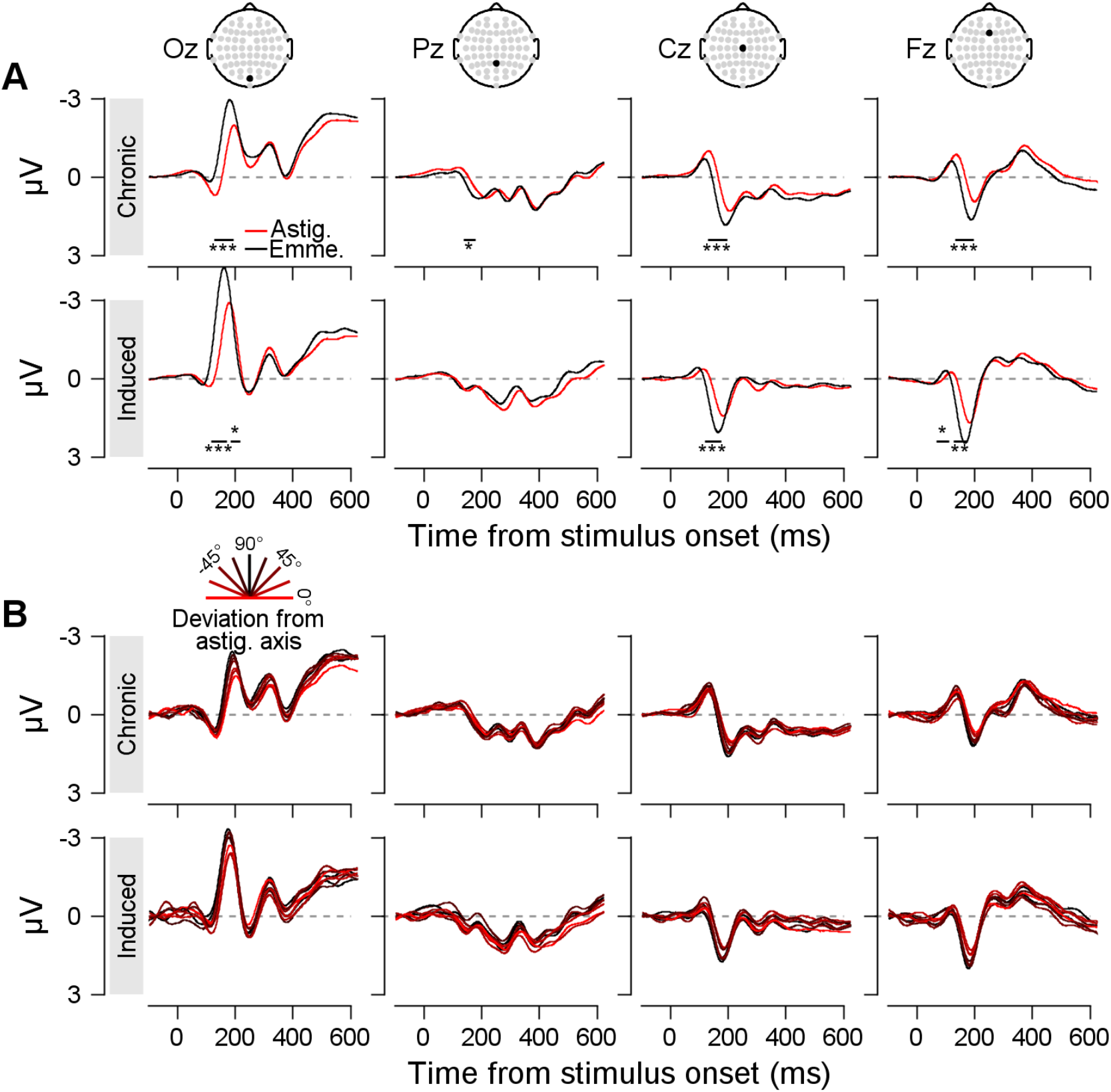
Event-related potential (ERP) results. **(A)** Grand-average ERPs in response to the eight Gabor patch orientations under astigmatic (red) and emmetropic (black) vision conditions are shown for both the chronic (top row) and induced astigmatism groups (bottom row) across electrodes Oz, Pz, Cz, and Fz. In the chronic astigmatism group, significant differences in ERPs between astigmatic and emmetropic vision conditions were observed at Oz electrode from 126 to 193 ms after stimulus onset, at Pz from 136 to 178 ms, at Cz from 129 to 198 ms, and at Fz from 133 to 198 ms. In the induced astigmatism group, significant differences in ERPs were found at Oz electrode from 116 to 172 ms and 184 to 218 ms after stimulus onset, at Cz 120 to 177 ms, and at Fz from 68 to 112 ms and 126 to 175 ms. Horizontal lines below the ERPs indicate significant time intervals, with stars denoting statistical significance levels (cluster-based permutation test; *, p < 0.05; **, p < 0.01; ***, p < 0.001). **(B)** ERPs under astigmatic vision conditions in response to Gabor stimuli tilted along the astigmatic axis (red) and in the orthogonal direction (black) are shown for both the chronic (upper row) and induced astigmatism groups (lower row) across electrodes Oz, Pz, Cz, and Fz. No significant interactions between orientations were observed in the ERPs (one-way ANOVA, ps > 0.05 for all time points).

**Supplementary Figure 4.**
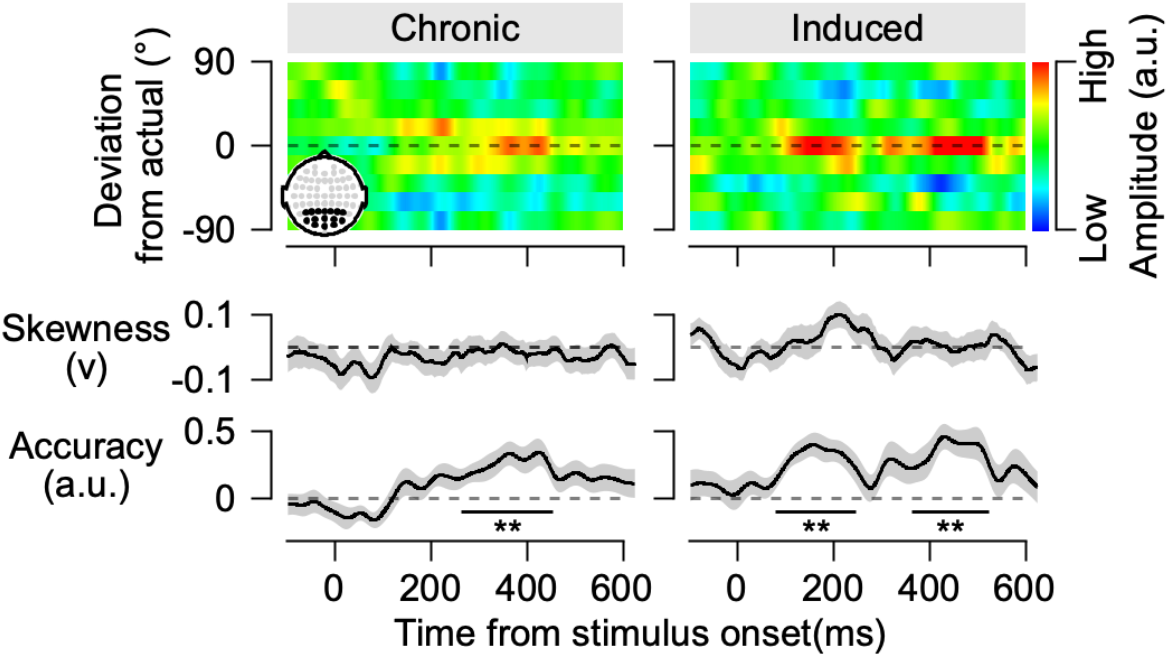
Orientation tuning responses in emmetropic vision condition. After full correction of the cylindrical refractive error, the orientation tuning responses for stimulus with oblique orientations were measured in the chronic (left) and induced (right) groups. The subfigures below indicate the skewness and strength of orientation information in the tuning responses. All notations are identical to those in Figure 2D.

**Supplementary Figure 5.**
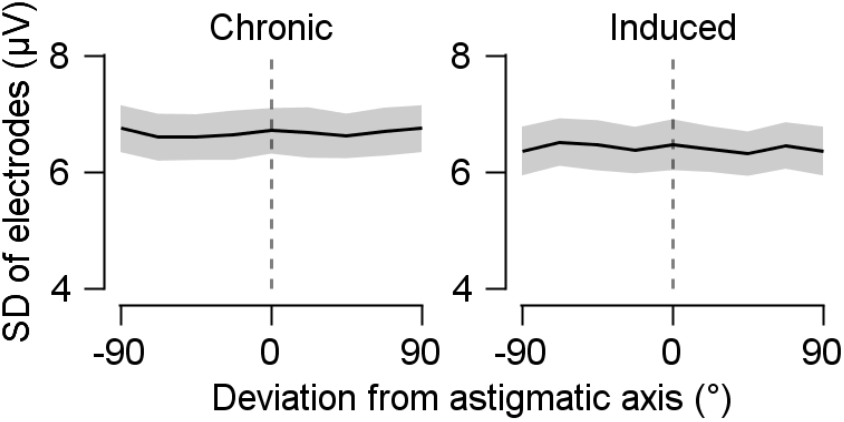
Stable variability of EEG activity across stimulus orientation. For each stimulus orientation, trial-by-trial variability (standard deviation, SD) was measured at posterior electrodes from 250 to 350 ms after stimulus onset. The SD was then averaged across time and electrodes, resulting in eight SD values per participant. These values were realigned so that SDs corresponding to the astigmatic axis were centered. Shaded regions represent ±1 SEM.

**Supplementary Figure 6.**
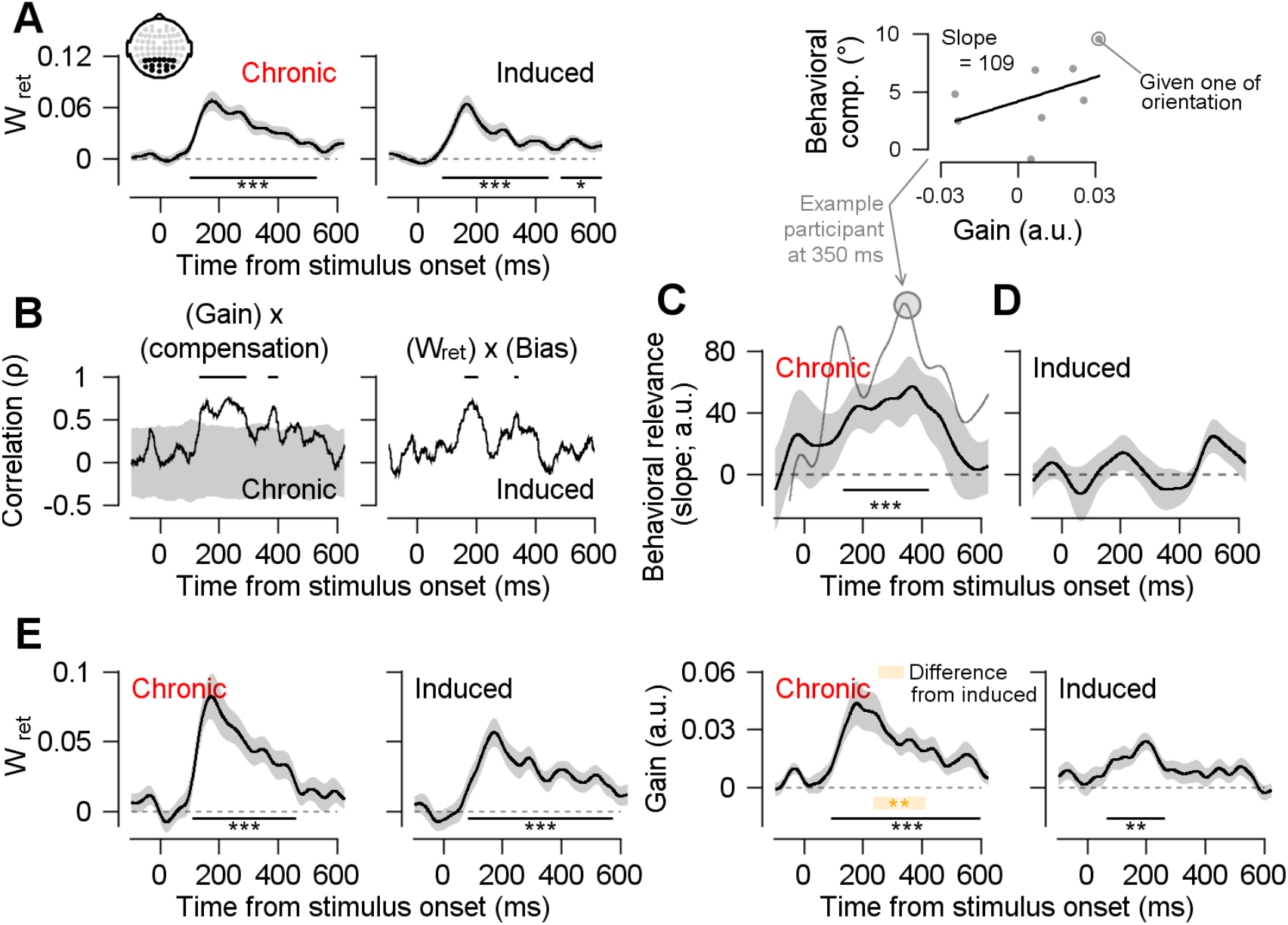
Distortion in retinal input and neural gain modulation. **(A)** Estimated weight of the retinal input component (in Figure 3A, left) in the orientation tuning responses (in Figure 2D). **(B)** Spearman’s correlation coefficient illustrating the relationship between neural gain modulation and perceptual compensation across individual participants in the chronic astigmatism group (left), and the relationship between retinal input weight and perceptual bias across individual participants in the induced astigmatism group (right). The shaded area represents the 95% confidence interval of the null distribution, generated from correlation coefficients between the neural gain modulations and behavioral reports when the mapping between the two was randomly shuffled. **(C and D)** Relationship between the strength of gain modulation and the reduction in perceptual bias (behavioral compensation). At each time point, a regression slope was computed to determine whether gain modulation across eight stimulus orientations co-varied with subsequent behavioral compensation within participants. A thin gray line represents an example subject, while an example regression result is illustrated using data points at 350 ms relative to stimulus onset in the top subfigure. **(E)** Simultaneous estimation of weights on the retinal input and the gain modulation components in the orientation tuning response. The overall results are consistent with those presented in the main text. All notations are identical to those in Figure 3B.

**Supplementary Figure 7.**
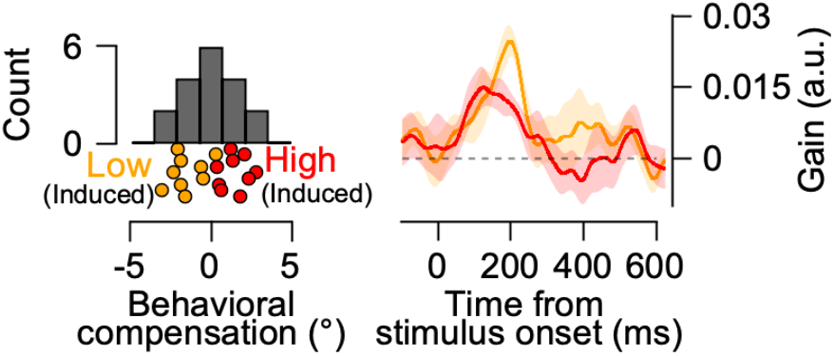
The gain modulation in the induced astigmatism group according to the degree of behavioral compensation. There were no significant differences in gain modulation when participants in the induced astigmatism group were divided into high and low behavioral compensation groups using the same method applied to the chronic astigmatism group (Figure 3C). All notations are identical to those in Figure 3C.

**Supplementary Figure 8.**
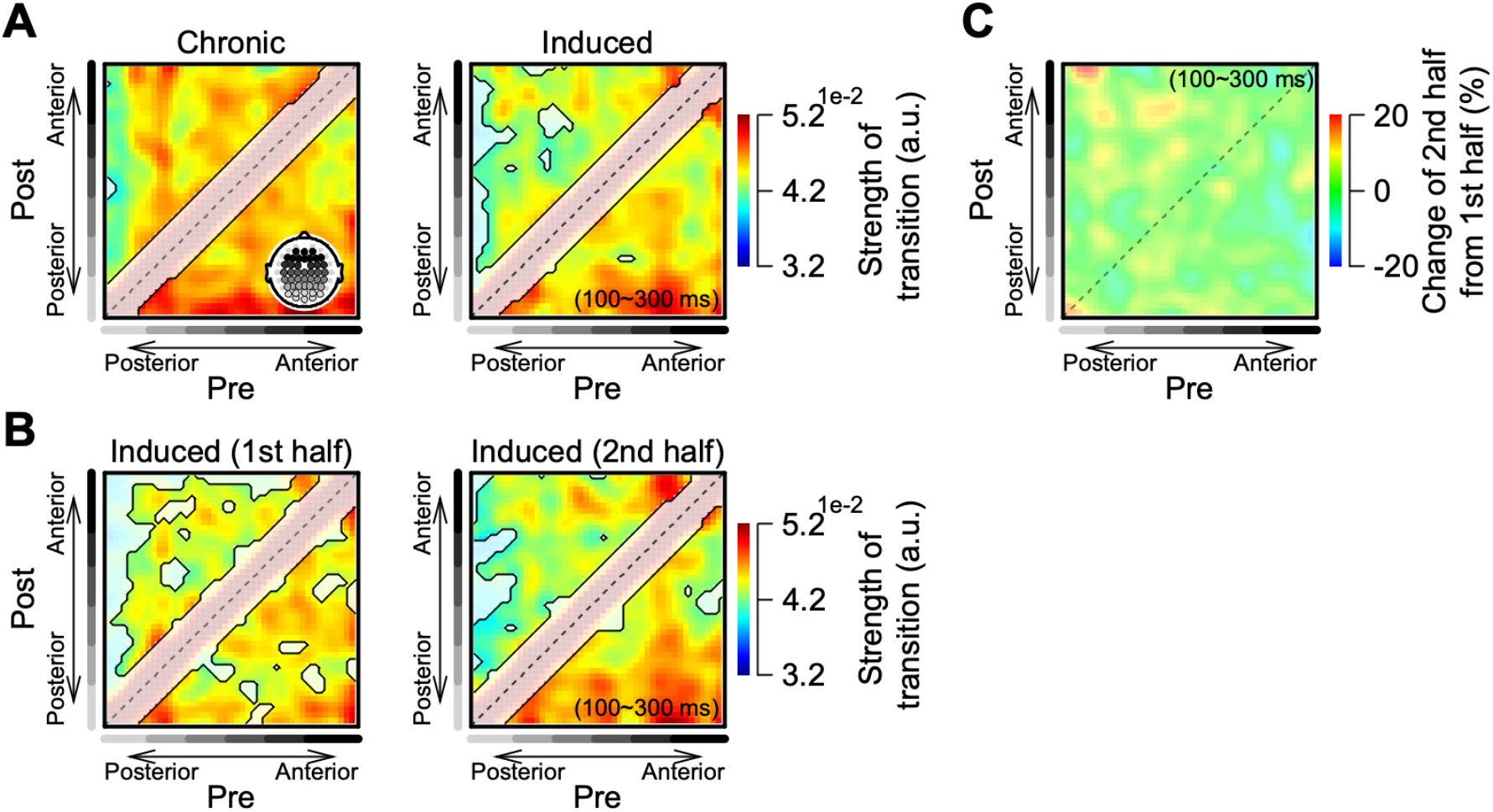
Transference of gain modulation in the chronic and induced astigmatism groups. **(A-B)** Absolute values of the push-pull gain modulation transition matrix for the chronic and induced astigmatism groups (A) and for the first and second halves of the experiment in the induced astigmatism group (B). Significant values are indicated by contours if they exceed the 95% confidence interval obtained from the null distribution of the transition matrix, which was generated from randomly shuffled gain modulation data. **(C)** Percentage change of the gain modulation transition matrix from the first half to the second half of the experiment in the chronic astigmatism group. All other notations are identical to those in Figure 4B.

**Supplementary Figure 9.**
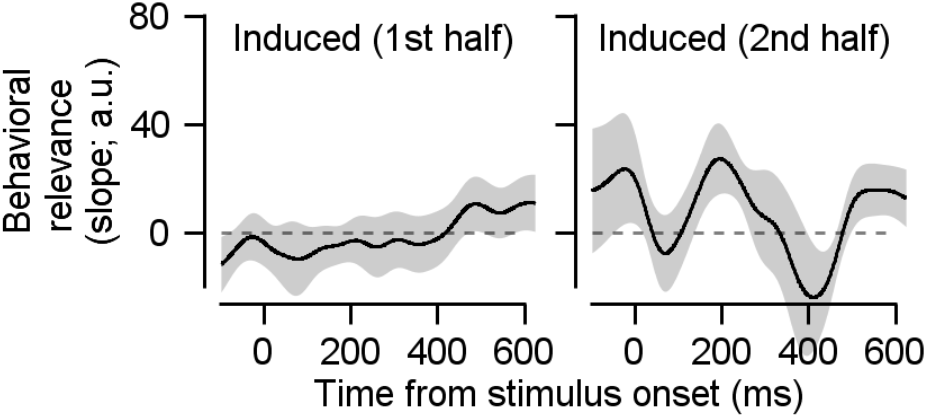
Relationship between gain modulation and behavioral compensation in the induced astigmatism group during the first and second halves of the experiment. As with Figure 3-1C-D, a regression slope was calculated to evaluate whether gain modulation co-flucuates with subsequent behavioral compensation. The shaded regions represent ±1 SEM.

## REFERENCES

Ajina, S., Jünemann, K., Sahraie, A., & Bridge, H. (2021). Increased Visual Sensitivity and Occipital Activity in Patients With Hemianopia Following Vision Rehabilitation. The Journal of Neuroscience: The Official Journal of the Society for Neuroscience, 41(28), 5994–6005. 10.1523/JNEUROSCI.2790-20.2021

Armstrong, K. M., Fitzgerald, J. K., & Moore, T. (2006). Changes in visual receptive fields with microstimulation of frontal cortex. Neuron, 50(5), 791–798. 10.1016/j.neuron.2006.05.010

Atchison, D. A., & Mathur, A. (2011). Visual acuity with astigmatic blur. Optometry and Vision Science: Official Publication of the American Academy of Optometry, 88(7), E798–805. 10.1097/OPX.0b013e3182186bc4

Baker, C. I., Peli, E., Knouf, N., & Kanwisher, N. G. (2005). Reorganization of visual processing in macular degeneration. The Journal of Neuroscience: The Official Journal of the Society for Neuroscience, 25(3), 614–618. 10.1523/JNEUROSCI.3476-04.2005

Bao, M., & Engel, S. A. (2012). Distinct mechanism for long-term contrast adaptation. Proceedings of the National Academy of Sciences of the United States of America, 109(15), 5898–5903. 10.1073/pnas.1113503109

Barbot, A., Das, A., Melnick, M. D., Cavanaugh, M. R., Merriam, E. P., Heeger, D. J., & Huxlin, K. R. (2021). Spared perilesional V1 activity underlies training-induced recovery of luminance detection sensitivity in cortically-blind patients. Nature Communications, 12(1), 6102. 10.1038/s41467-021-26345-1

Barbot, A., Park, W. J., Ng, C. J., Zhang, R.-Y., Huxlin, K. R., Tadin, D., & Yoon, G. (2021). Functional reallocation of sensory processing resources caused by long-term neural adaptation to altered optics. ELife, 10. 10.7554/eLife.58734

Bavelier, D., Levi, D. M., Li, R. W., Dan, Y., & Hensch, T. K. (2010). Removing brakes on adult brain plasticity: from molecular to behavioral interventions. The Journal of Neuroscience: The Official Journal of the Society for Neuroscience, 30(45), 14964–14971. 10.1523/JNEUROSCI.4812-10.2010

Benavente-Pérez, A., Nour, A., & Troilo, D. (2014). Axial eye growth and refractive error development can be modified by exposing the peripheral retina to relative myopic or hyperopic defocus. Investigative Ophthalmology & Visual Science, 55(10), 6765–6773. 10.1167/iovs.14-14524

Benjamin, W. J., & Borish, I. M. (2006). Borish’s Clinical Refraction (2nd ed.). Butterworth-Heinemann. https://play.google.com/store/books/details?id=Yeq8QAAACAAJ

Berens, P. (2009). CircStat: A MATLAB Toolbox for Circular Statistics. Journal of Statistical Software, 31, 1–21. 10.18637/jss.v031.i10

Beyeler, M., Rokem, A., Boynton, G. M., & Fine, I. (2017). Learning to see again: biological constraints on cortical plasticity and the implications for sight restoration technologies. Journal of Neural Engineering, 14(5), 051003. 10.1088/1741-2552/aa795e

Bigdely-Shamlo, N., Mullen, T., Kothe, C., Su, K.-M., & Robbins, K. A. (2015). The PREP pipeline: standardized preprocessing for large-scale EEG analysis. Frontiers in Neuroinformatics, 9, 16. 10.3389/fninf.2015.00016

Bloem, I. M., & Ling, S. (2019). Normalization governs attentional modulation within human visual cortex. Nature Communications, 10(1), 5660. 10.1038/s41467-019-13597-1

Brainard, D. H. (1997). The Psychophysics Toolbox. Spatial Vision, 10(4), 433–436. https://www.ncbi.nlm.nih.gov/pubmed/9176952

Byers, A., & Serences, J. T. (2014). Enhanced attentional gain as a mechanism for generalized perceptual learning in human visual cortex. Journal of Neurophysiology, 112(5), 1217–1227. 10.1152/jn.00353.2014

Castaldi, E., Lunghi, C., & Morrone, M. C. (2020). Neuroplasticity in adult human visual cortex. Neuroscience and Biobehavioral Reviews, 112, 542–552. 10.1016/j.neubiorev.2020.02.028

Cavanaugh, M. R., Barbot, A., Carrasco, M., & Huxlin, K. R. (2019). Feature-based attention potentiates recovery of fine direction discrimination in cortically blind patients. Neuropsychologia, 128, 315–324. 10.1016/j.neuropsychologia.2017.12.010

Chance, F. S., Abbott, L. F., & Reyes, A. D. (2002). Gain modulation from background synaptic input. Neuron, 35(4), 773–782. 10.1016/s0896-6273(02)00820-6

Cottaris, N. P., Jiang, H., Ding, X., Wandell, B. A., & Brainard, D. H. (2019). A computational-observer model of spatial contrast sensitivity: Effects of wave-front-based optics, cone-mosaic structure, and inference engine. Journal of Vision, 19(4), 8. 10.1167/19.4.8

Dosher, B., & Lu, Z.-L. (2017). Visual Perceptual Learning and Models. Annual Review of Vision Science, 3, 343–363. 10.1146/annurev-vision-102016-061249

Fellous, J.-M., Rudolph, M., Destexhe, A., & Sejnowski, T. J. (2003). Synaptic background noise controls the input/output characteristics of single cells in an in vitro model of in vivo activity. Neuroscience, 122(3), 811–829. 10.1016/j.neuroscience.2003.08.027

Georgeson, M. A., & Sullivan, G. D. (1975). Contrast constancy: deblurring in human vision by spatial frequency channels. The Journal of Physiology, 252(3), 627–656. 10.1113/jphysiol.1975.sp011162

Haak, K. V., Fast, E., Bao, M., Lee, M., & Engel, S. A. (2014). Four days of visual contrast deprivation reveals limits of neuronal adaptation. Current Biology: CB, 24(21), 2575–2579. 10.1016/j.cub.2014.09.027

Hajonides, J. E., Nobre, A. C., van Ede, F., & Stokes, M. G. (2021). Decoding visual colour from scalp electroencephalography measurements. NeuroImage, 237, 118030. 10.1016/j.neuroimage.2021.118030

Hashemi, H., Fotouhi, A., Yekta, A., Pakzad, R., Ostadimoghaddam, H., & Khabazkhoob, M. (2018). Global and regional estimates of prevalence of refractive errors: Systematic review and meta-analysis. Journal of Current Ophthalmology, 30(1), 3–22. 10.1016/j.joco.2017.08.009

Hubel, D. H., & Wiesel, T. N. (1959). Receptive fields of single neurones in the cat’s striate cortex. The Journal of Physiology, 148(3), 574–591. 10.1113/jphysiol.1959.sp006308

Jehee, J. F. M., Ling, S., Swisher, J. D., van Bergen, R. S., & Tong, F. (2012). Perceptual learning selectively refines orientation representations in early visual cortex. The Journal of Neuroscience: The Official Journal of the Society for Neuroscience, 32(47), 16747–53a. 10.1523/JNEUROSCI.6112-11.2012

Jing, R., Yang, C., Huang, X., & Li, W. (2021). Perceptual learning as a result of concerted changes in prefrontal and visual cortex. Current Biology: CB, 31(20), 4521–4533.e3. 10.1016/j.cub.2021.08.007

Jones, J. P., & Palmer, L. A. (1987). An evaluation of the two-dimensional Gabor filter model of simple receptive fields in cat striate cortex. Journal of Neurophysiology, 58(6), 1233–1258. 10.1152/jn.1987.58.6.1233

Karni, A., & Sagi, D. (1991). Where practice makes perfect in texture discrimination: evidence for primary visual cortex plasticity. Proceedings of the National Academy of Sciences of the United States of America, 88(11), 4966–4970. 10.1073/pnas.88.11.4966

Kim, T.-I., Alió Del Barrio, J. L., Wilkins, M., Cochener, B., & Ang, M. (2019). Refractive surgery. The Lancet, 393(10185), 2085–2098. 10.1016/S0140-6736(18)33209-4

Kobashi, H., Kamiya, K., Shimizu, K., Kawamorita, T., & Uozato, H. (2012). Effect of axis orientation on visual performance in astigmatic eyes. Journal of Cataract and Refractive Surgery, 38(8), 1352–1359. 10.1016/j.jcrs.2012.03.032

Kriegeskorte, N., Goebel, R., & Bandettini, P. (2006). Information-based functional brain mapping. Proceedings of the National Academy of Sciences of the United States of America, 103(10), 3863–3868. 10.1073/pnas.0600244103

Lazzouni, L., & Lepore, F. (2014). Compensatory plasticity: time matters. Frontiers in Human Neuroscience, 8, 340. 10.3389/fnhum.2014.00340

Lee, T. S. (1996). Image representation using 2D Gabor wavelets. IEEE Transactions on Pattern Analysis and Machine Intelligence, 18(10), 959–971. 10.1109/34.541406

Li, W. (2016). Perceptual Learning: Use-Dependent Cortical Plasticity. Annual Review of Vision Science, 2, 109–130. 10.1146/annurev-vision-111815-114351

Liu, Y., Yu, C., Liang, M., Li, J., Tian, L., Zhou, Y., Qin, W., Li, K., & Jiang, T. (2007). Whole brain functional connectivity in the early blind. Brain: A Journal of Neurology, 130(Pt 8), 2085–2096. 10.1093/brain/awm121

Makeig, S., Bell, A., Jung, T.-P., & Sejnowski, T. J. (1995). Independent component analysis of electroencephalographic data. Advances in Neural Information Processing Systems, 8. https://proceedings.neurips.cc/paper/1995/hash/754dda4b1ba34c6fa89716b85d68532b-Abstract.html

Mante, V., Sussillo, D., Shenoy, K. V., & Newsome, W. T. (2013). Context-dependent computation by recurrent dynamics in prefrontal cortex. Nature, 503(7474), 78–84. 10.1038/nature12742

Maris, E., & Oostenveld, R. (2007). Nonparametric statistical testing of EEG- and MEG-data. Journal of Neuroscience Methods, 164(1), 177–190. 10.1016/j.jneumeth.2007.03.024

McLean, R. C., & Wallman, J. (2003). Severe astigmatic blur does not interfere with spectacle lens compensation. Investigative Ophthalmology & Visual Science, 44(2), 449–457. 10.1167/iovs.01-0670

Mitchell, D. E., Freeman, R. D., Millodot, M., & Haegerstrom, G. (1973). Meridional amblyopia: evidence for modification of the human visual system by early visual experience. Vision Research, 13(3), 535–558. 10.1016/0042-6989(73)90023-0

Mognon, A., Jovicich, J., Bruzzone, L., & Buiatti, M. (2011). ADJUST: An automatic EEG artifact detector based on the joint use of spatial and temporal features. Psychophysiology, 48(2), 229–240. 10.1111/j.1469-8986.2010.01061.x

Moldakarimov, S., Bazhenov, M., & Sejnowski, T. J. (2014). Top-down inputs enhance orientation selectivity in neurons of the primary visual cortex during perceptual learning. PLoS Computational Biology, 10(8), e1003770. 10.1371/journal.pcbi.1003770

Moore, T., & Armstrong, K. M. (2003). Selective gating of visual signals by microstimulation of frontal cortex. Nature, 421(6921), 370–373. 10.1038/nature01341

Mullen, T. R., Kothe, C. A. E., Chi, Y. M., Ojeda, A., Kerth, T., Makeig, S., Jung, T.-P., & Cauwenberghs, G. (2015). Real-Time Neuroimaging and Cognitive Monitoring Using Wearable Dry EEG. IEEE Transactions on Bio-Medical Engineering, 62(11), 2553–2567. 10.1109/TBME.2015.2481482

Murphy, B. K., & Miller, K. D. (2009). Balanced amplification: a new mechanism of selective amplification of neural activity patterns. Neuron, 61(4), 635–648. 10.1016/j.neuron.2009.02.005

Myers, N. E., Rohenkohl, G., Wyart, V., Woolrich, M. W., Nobre, A. C., & Stokes, M. G. (2015). Testing sensory evidence against mnemonic templates. ELife, 4, e09000. 10.7554/eLife.09000

Ng, C. J., Blake, R., Banks, M. S., Tadin, D., & Yoon, G. (2021). Optics and neural adaptation jointly limit human stereovision. Proceedings of the National Academy of Sciences of the United States of America, 118(23), e2100126118. 10.1073/pnas.2100126118

Nienborg, H., Cohen, M. R., & Cumming, B. G. (2012). Decision-related activity in sensory neurons: correlations among neurons and with behavior. Annual Review of Neuroscience, 35(1), 463–483. 10.1146/annurev-neuro-062111-150403

Ohlendorf, A., Tabernero, J., & Schaeffel, F. (2011). Neuronal adaptation to simulated and optically-induced astigmatic defocus. Vision Research, 51(6), 529–534. 10.1016/j.visres.2011.01.010

Olofsson, P. (2012). Probability, Statistics, and Stochastic Processes. Wiley & Sons, Limited, John. https://openlibrary.org/books/OL33466428M/Probability_Statistics_and_Stochastic_Processes

Pelli, D. G. (1997). The VideoToolbox software for visual psychophysics: transforming numbers into movies. Spatial Vision, 10(4), 437–442. https://www.ncbi.nlm.nih.gov/pubmed/9176953

Portilla, J., & Simoncelli, E. P. (2000). A Parametric Texture Model Based on Joint Statistics of Complex Wavelet Coefficients. International Journal of Computer Vision, 40(1), 49–70. 10.1023/A:1026553619983

Purves, D., & White, L. E. (1994). Monocular preferences in binocular viewing. Proceedings of the National Academy of Sciences of the United States of America, 91(18), 8339–8342. 10.1073/pnas.91.18.8339

Radhakrishnan, A., Dorronsoro, C., Sawides, L., Webster, M. A., & Marcos, S. (2015). A cyclopean neural mechanism compensating for optical differences between the eyes. Current Biology: CB, 25(5), R188–9. 10.1016/j.cub.2015.01.027

Regan, D. (1973). Rapid objective refraction using evoked brain potentials. Investigative Ophthalmology & Visual Science, 12(9), 669–679. https://iovs.arvojournals.org/article.aspx?articleid=2203269

Sanayei, M., Chen, X., Chicharro, D., Distler, C., Panzeri, S., & Thiele, A. (2018). Perceptual learning of fine contrast discrimination changes neuronal tuning and population coding in macaque V4. Nature Communications, 9(1), 4238. 10.1038/s41467-018-06698-w

Sawides, L., Marcos, S., Ravikumar, S., Thibos, L., Bradley, A., & Webster, M. (2010). Adaptation to astigmatic blur. Journal of Vision, 10(12), 22. 10.1167/10.12.22

Schoups, A., Vogels, R., Qian, N., & Orban, G. (2001). Practising orientation identification improves orientation coding in V1 neurons. Nature, 412(6846), 549–553. 10.1038/35087601

Sokol, S. (1983). Abnormal evoked potential latencies in amblyopia. The British Journal of Ophthalmology, 67(5), 310–314. 10.1136/bjo.67.5.310

Son, S., Moon, J., Kang, H., Kim, Y.-J., & Lee, J. (2021). Induced astigmatism biases the orientation information represented in multivariate electroencephalogram activities. Human Brain Mapping, 42(13), 4336–4347. 10.1002/hbm.25550

Son, S., Shim, W. M., Kang, H., & Lee, J. (2022). Automatic compensation enhances the orientation perception in chronic astigmatism. Scientific Reports, 12(1), 3710. 10.1038/s41598-022-07788-y

Villegas, E. A., Alcón, E., & Artal, P. (2014). Minimum amount of astigmatism that should be corrected. Journal of Cataract and Refractive Surgery, 40(1), 13–19. 10.1016/j.jcrs.2013.09.010

Vinas, M., Sawides, L., de Gracia, P., & Marcos, S. (2012). Perceptual adaptation to the correction of natural astigmatism. PloS One, 7(9), e46361. 10.1371/journal.pone.0046361

Watanabe, T., & Sasaki, Y. (2015). Perceptual learning: toward a comprehensive theory. Annual Review of Psychology, 66, 197–221. 10.1146/annurev-psych-010814-015214

Webster, M. A., Georgeson, M. A., & Webster, S. M. (2002). Neural adjustments to image blur. Nature Neuroscience, 5(9), 839–840. 10.1038/nn906

Williams, K. M., Verhoeven, V. J. M., Cumberland, P., Bertelsen, G., Wolfram, C., Buitendijk, G. H. S., Hofman, A., van Duijn, C. M., Vingerling, J. R., Kuijpers, R. W. A. M., Höhn, R., Mirshahi, A., Khawaja, A. P., Luben, R. N., Erke, M. G., von Hanno, T., Mahroo, O., Hogg, R., Gieger, C., … Hammond, C. J. (2015). Prevalence of refractive error in Europe: the European Eye Epidemiology (E(3)) Consortium. European Journal of Epidemiology, 30(4), 305–315. 10.1007/s10654-015-0010-0

Wolff, M. J., Ding, J., Myers, N. E., & Stokes, M. G. (2015). Revealing hidden states in visual working memory using electroencephalography. Frontiers in Systems Neuroscience, 9(September), 123. 10.3389/fnsys.2015.00123

Wolff, M. J., Jochim, J., Akyürek, E. G., & Stokes, M. G. (2017). Dynamic hidden states underlying working-memory-guided behavior. Nature Neuroscience, 20(6), 864–871. 10.1038/nn.4546

Wolffsohn, J. S., Bhogal, G., & Shah, S. (2011). Effect of uncorrected astigmatism on vision. Journal of Cataract and Refractive Surgery, 37(3), 454–460. 10.1016/j.jcrs.2010.09.022

Wong, T. Y., Foster, P. J., Hee, J., Ng, T. P., Tielsch, J. M., Chew, S. J., Johnson, G. J., & Seah, S. K. (2000). Prevalence and risk factors for refractive errors in adult Chinese in Singapore. Investigative Ophthalmology & Visual Science, 41(9), 2486–2494. https://www.ncbi.nlm.nih.gov/pubmed/10937558

Yan, Y., Rasch, M. J., Chen, M., Xiang, X., Huang, M., Wu, S., & Li, W. (2014). Perceptual training continuously refines neuronal population codes in primary visual cortex. Nature Neuroscience, 17(10), 1380–1387. 10.1038/nn.3805

Yang, T., & Maunsell, J. H. R. (2004). The effect of perceptual learning on neuronal responses in monkey visual area V4. The Journal of Neuroscience: The Official Journal of the Society for Neuroscience, 24(7), 1617–1626. 10.1523/JNEUROSCI.4442-03.2004

Yehezkel, O., Sagi, D., Sterkin, A., Belkin, M., & Polat, U. (2010). Learning to adapt: Dynamics of readaptation to geometrical distortions. Vision Research, 50(16), 1550–1558. 10.1016/j.visres.2010.05.014

Zhang, P., Bao, M., Kwon, M., He, S., & Engel, S. A. (2009). Effects of orientation-specific visual deprivation induced with altered reality. Current Biology: CB, 19(22), 1956–1960. 10.1016/j.cub.2009.10.018

